# Highly replicated evolution of parapatric ecotypes

**DOI:** 10.1101/2020.02.05.936401

**Authors:** Maddie E. James, Henry Arenas-Castro, Jeffrey S. Groh, Scott L. Allen, Jan Engelstädter, Daniel Ortiz-Barrientos

## Abstract

Parallel evolution of ecotypes occurs when selection independently drives the evolution of similar traits across similar environments. The multiple origin of ecotypes is often inferred on the basis of a phylogeny which clusters populations according to geographic location and not by the environment they occupy. However, the use of phylogenies to infer parallel evolution in closely related populations is problematic because gene flow and incomplete lineage sorting can uncouple the genetic structure at neutral markers from the colonization history of populations. Here, we demonstrate multiple origins within ecotypes of an Australian wildflower, *Senecio lautus*. We observed strong genetic structure as well as phylogenetic clustering by geography and show that this is unlikely due to gene flow between parapatric ecotypes, which is surprisingly low. We further confirm this analytically by demonstrating that phylogenetic distortion due to gene flow often requires higher levels of migration than those observed in *S. lautus*. Our results imply that selection can repeatedly create similar phenotypes despite the perceived homogenizing effects of gene flow.

## Introduction

Parallel evolution occurs when populations evolve similar traits after repeatedly and independently colonizing similar habitats (Schluter and Nagel 1995). When the distribution of habitats is patchy, phenotypically similar populations frequently occur next to other contrasting phenotypes (e.g., plant species adapted to serpentine and non-serpentine soils in Scandinavia; Berglund et al. 2003, and marine snails adapted to crab predators or wave action along the rocky coasts of Europe; Johannesson et al. 2010). Parallel evolution by natural selection creates consistent patterns of phenotypic similarity and divergence that can extend to morphological (Elmer et al. 2010; Ravinet et al. 2013; Perreault-Payette et al. 2017), behavioral (York and Fernald 2017), and reproductive (Smith and Rausher 2011) traits, yet there is often variation in the extent to which replicate populations exhibit similar phenotypes (see Bolnick et al. 2018). The nature of parallel trait evolution largely depends on the demographic history of the system under investigation, where the interplay of geography, gene flow, and natural selection along with the genetic architecture of traits determines its repeatability (Orr 2005; Stern and Orgogozo 2009; Rosenblum et al. 2014; Lenormand et al. 2016; Stoltzfus and McCandlish 2017; Blount et al. 2018; Yeaman et al. 2018). However, it is surprisingly rare for studies of parallel evolution to convincingly demonstrate that populations exhibiting similar phenotypes have arisen in an independent and repeated fashion (‘multiple origin’ scenario). Ruling out alternative demographic scenarios, such as a single origin of ecotypes followed by gene flow upon secondary contact, is seldom performed (but see work in *Littorina* and stickleback, Quesada et al. 2007; Bierne et al. 2013; Butlin et al. 2014; Pérez-Pereira et al. 2017, and see Ostevik et al. 2012 for a critical review of the evidence in plants). In light of this, researchers may incorrectly assume a parallel demographic history, leading to inaccurate inferences about the prevalence of parallel evolution in nature.

Typically, researchers identify parallel evolution by natural selection by asking whether phylogenetic clustering of populations (estimated in the absence of gene flow) coincides with the geographic distribution of populations (geography) and not with the habitat in which they occur (ecology) (Allender et al. 2003; Quesada et al. 2007; Johannesson et al. 2010; Butlin et al. 2014; Trucchi et al. 2017). This is because genetic clustering of geographically close populations implies dispersal might be geographically restricted (i.e., isolation by distance; Wright 1943), and colonization of contrasting and neighboring habitats might have occurred independently many times. The rationale for this argument is that the genome-wide phylogenetic history across the geographic range of populations can reveal the history of adaptation. That is, if the repeated evolution of similar phenotypes appears to have taken place on different genetic backgrounds (lineages), then the genetic changes that drove adaptation likely occurred independently. By genetic changes, we refer specifically to independent allele frequency changes driven by similar natural selection pressures.

The above argument rests upon the assumption that the genome-wide pattern of relatedness accurately depicts the history of the loci underlying adaptation, although this is not necessarily the case. For example, alternative historical scenarios could also lead to clustering of populations by geography, and must be ruled out before examining the evolution of traits in light of parallel evolution (Endler 1977; Barton and Hewitt 1985; Coyne and Orr 2004; Bierne et al. 2013). For instance, a recent phylogenomic analysis of *Mimulus* section *Erythranthe* overturned a classical example of putative convergence in hummingbird pollination (Beardsley et al. 2004; Freeman and Herron 2004). The authors discovered that hummingbird pollination in the genus arose only once, yet over a large portion of the genome, sympatric populations of *M. cardinalis* (hummingbird-pollinated) and *M. lewisii* (bee-pollinated) appear as sister taxa in the phylogeny because of multiple introgression events after periods of allopatry (Nelson et al. 2021). To understand how demography, geography and selection contribute to our interpretations of parallel evolution, first consider a scenario where an ancestral population gives rise to two locally adapted populations that occupy ecologically distinct yet geographically proximate habitats (hereafter ecotypes, Fig. 1A). These two populations repeatedly migrate to new localities, where each time the same contrasting habitats are geographically close. This scenario of a single split followed by range expansion of two ecotypes does not involve a parallel adaptation history because each ecotype only arose once (rather than multiple independent times after independent colonization of contrasting habitats). Because gene flow is either not possible after the original ecotypic split due to complete reproductive isolation, or gene flow does not homogenize adjacent populations after range expansion, populations sharing the same ecology form reciprocally monophyletic clades in a phylogeny (Fig. 1A, *Single origin without gene flow*).

**Fig 1.**
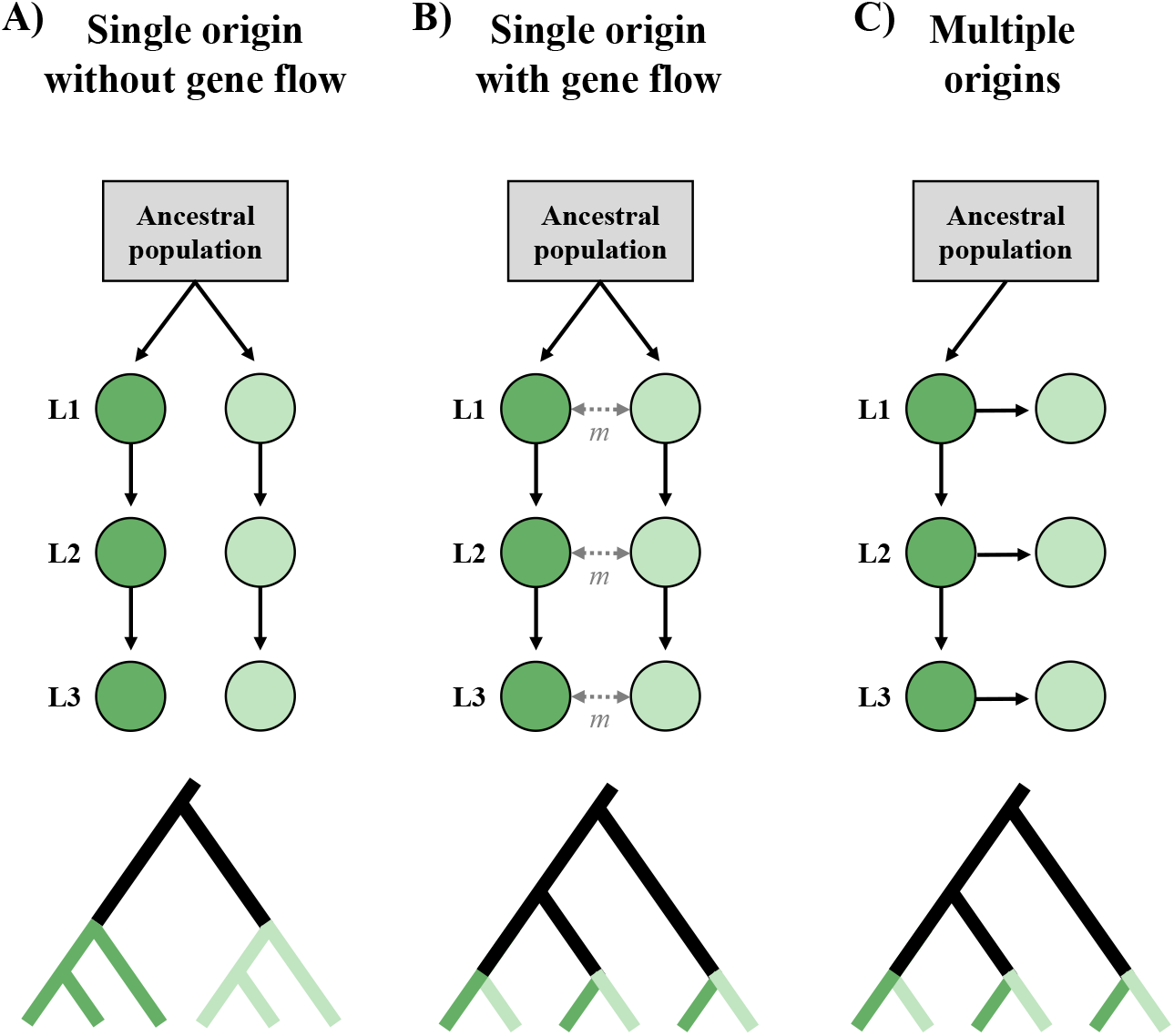
The colonization history and phylogenetic topology for alternate origin scenarios. Schematic diagram representing the colonization history and phylogenetic topology of two ecotypes (dark green and light green) from an ancestral population (grey) for three origin scenarios. Solid arrows depict the sequence of colonization. Double headed dotted arrows represent gene flow (*m*) between the ecotypes within each locality. L1, L2 and L3 represent three geographically distant localities, where a population of each ecotype resides. **(A)** Within a single origin scenario, the two ecotypes arise once from the ancestor, followed by range expansion. In the absence of gene flow, ecotypes form monophyletic clades within the phylogeny. **(B)** The single origin with gene flow scenario involves gene flow upon secondary contact between the ecotypes within each locality. Here, the observed phylogenetic topology shows populations clustering according to their geographic distribution. **(C)** Within a multiple origin scenario, the ancestral (dark green) ecotype arises once from the ancestor followed by range expansion, with the derived (light green) populations independently arising from each dark green population. Populations phylogenetically cluster according to their geographic distribution, which can be indistinguishable from a single origin with gene flow scenario **(B)**.

Nevertheless, if there is sufficient gene flow between geographically close populations from two ecotypes that originated only once (like the *Mimulus* example described above), the original phylogenetic signal of reciprocal monophyly could be eroded (Endler 1977; Barton and Hewitt 1985; Coyne and Orr 2004; Bierne et al. 2013). In other words, as the original genome-wide phylogenetic signal of a single origin disappears, populations become most related to their neighboring population and not to the other populations of the same ecotype. Therefore, gene flow can result in grouping of populations by geography rather than ecology (Fig. 1B, *Single origin with gene flow*). This phylogenetic signal is identical to that of true parallel evolution (a multiple origin scenario), where the derived ecotype arises multiple independent times from the ancestral ecotype (Fig. 1C, *Multiple origins*). Gene flow dynamics can thus fundamentally alter our interpretation of parallel evolution, to the extent that we can mistakenly infer parallel evolution in systems where secondary contact after range expansion of a single origin has obscured the history of locally adapted populations (Endler 1977; Barton and Hewitt 1985; Coyne and Orr 2004; Bierne et al. 2013).

Despite the potential for gene flow to paint a false picture of the phylogenetic history of multiple populations adapted to the same environment, it is important to note that non-monophyly of populations from the same ecotype is not a requirement for parallel evolution in a more general sense. This is because it is possible that two populations of the same ecotype that are true phylogenetic sister groups did in fact adapt independently. However, non-monophyly of each ecotype is a requirement in systems where parallel evolution coincides with a patchy geographic distribution of populations (pairs of ecotypes in multiple localities; Bierne et al. 2013), where the phylogenetic line of reasoning is commonly employed.

In systems of parallel evolution, gene flow is frequently detected between populations, especially when contrasting ecotypes are in close geographic proximity (i.e., parapatry). Although not all levels of gene flow have the same effect on the genetic record of colonization history (Bierne et al. 2013), gene flow between ecotypes must be taken into account when demonstrating parallel origins within a system. However, only very few systems have thoroughly investigated the demographic history of populations, and even fewer have used coalescent modelling or simulations to address whether the estimated levels of gene flow could have obscured the observed phylogeny. A system that has clearly demonstrated the parallel origins of contrasting populations in the presence of gene flow is the marine snail *Littorina saxatilis*. Multiple lines of evidence suggest the wave and crab ecotypes evolved multiple independent times along rocky coastlines (Quesada et al. 2007; Johannesson et al. 2010; Bierne et al. 2013; Butlin et al. 2014; Pérez-Pereira et al. 2017). Other systems providing evidence for parallel evolution include Lake Victoria cichlids (Meier et al. 2017) and alpine and montane *Heliosperma pusillum* ecotypes (Trucchi et al. 2017). Also, an obvious case showing multiple origins is when parallel evolution occurs between geographically distant populations where gene flow could not have obscured the phylogenetic signal and demographic history of populations (e.g., threespine stickleback populations that colonized freshwater environments on separate continents; Magalhaes et al. 2020). However, in other systems, where gene flow is moderate between ecotypes (Rougemont et al. 2015; Le Moan et al. 2016; Rougeux et al. 2017; Herman et al. 2018; Rougeux et al. 2019), it remains unclear to what extent gene flow contributed to the signal of parallel evolution.

We must keep in mind that identifying the genetic basis of parallel trait evolution often provides unambiguous evidence for parallel evolution of ecotypes. For instance, in sticklebacks, the repeated evolution of the reduction of pelvic armor in separate populations occurred via different mutations in the same gene (Chan et al. 2010), suggesting this adaptive trait has arisen and been selected for multiple independent times. In contrast, in systems where the exact same mutation is repeatedly involved in adaptation (e.g., Colosimo et al. 2005), it is challenging to identify whether the adaptive mutation was repeatedly and independently selected for in each population (either from de-novo mutations or via standing genetic variation; Roesti et al. 2014; Lee and Coop 2019). Knowing the causal genes of adaptation is ideal as the demographic history of individual adaptive loci can be modelled, avoiding the complications of distinguishing between single and multiple origins using neutral polymorphisms (as described above). However, directly isolating the specific genes involved in adaptation is infeasible in most non-model organisms or when the genetic architecture of adaptation is highly polygenic (Tiffin and Ross-Ibarra 2014; Yeaman 2015).

The above considerations suggest that without knowing the genetic basis of parallel adaptation, we should carefully characterize and interpret the phylogeographic history to understand the level of independence in systems where populations are adapted to similar environments. Such an approach is necessary to demonstrate that natural selection has independently acted in separate populations during the repeated adaptation to similar environments. This knowledge paves the way for future research on dissecting the molecular basis of parallel adaptation, and has implications for our understanding of how repeatable and predictable evolution can be. In this work, we thoroughly characterize the phylogenetic and demographic history of *Senecio lautus*, an Australian wildflower that appears to have evolved multiple times in parapatry into two contrasting coastal forms called Dune and Headland ecotypes (Roda, Ambrose, et al. 2013; Melo et al. 2014). The two forms differ in their growth habit: the Dune ecotype is erect and inhabits sand dunes, and the Headland ecotype is prostrate, forming matts on the ground of rocky headlands (Ali 1964; Radford et al. 2004; Thompson 2005). These locally adapted populations (Melo et al. 2014; Richards et al. 2016; Richards and Ortiz-Barrientos 2016; Walter et al. 2016; Walter, Wilkinson, et al. 2018; Wilkinson et al. 2021) are separated by strong extrinsic reproductive isolation (Melo et al. 2014; Richards et al. 2016), and populations exhibit similar morphology within each ecotype (Walter, Aguirre, et al. 2018; James et al. 2020). With this work we hope to clearly illustrate how the demographic history of populations affects the evidence for the independent and repeated origins of parapatric ecotypes.

Previous work using pools of DNA sequences from multiple coastal, inland, alpine, and woodland *S. lautus* ecotypes found that strong isolation by distance separated all populations along the coast and that geography, not ecology, explained the phylogenetic clustering of coastal populations (Roda, Ambrose, et al. 2013). Although these results initially suggested that the Dune and Headland ecotypes evolved in parallel, it remains unclear if gene flow could be responsible for this pattern of ecotypic and geographic differentiation. Therefore, our inferences on the number of independent colonization and origin events of Dune and Headland populations remains unclear. Here, we directly estimate patterns of gene flow between 23 Dune and Headland populations, as well as other parameters important for characterizing the demographic history of this system. We create a coalescent model and undertake simulations to explore the conditions that could erode a signal of phylogenetic monophyly of each ecotype, thus enabling us to gain further confidence in our conclusions about parallel parapatric divergence in this system. Our results enhance our understanding of the nature of parallel evolution and pave the way for analyses of parallel trait evolution driven by natural selection in plants, where cases of parallelism remain understudied.

## Results

### Populations cluster by geography and not by ecology

To ask whether populations cluster according to their geographic distribution, we explored broad patterns of genetic clustering across the 23 Dune and Headland *S. lautus* populations (Fig. 2A). Phylogenetic inference in *IQ-TREE* provides evidence against a single origin scenario: neither ecotype forms a monophyletic clade, and parapatric Dune-Headland populations are often sister-taxa, consistent with a possible multiple origin of ecotypes (Fig. 2B). To visualize the major genetic structure within *fastSTRUCTURE*, we plotted the lowest K-values that capture the major structure in the data as suggested by Pritchard et al. (2000) and Lawson et al. (2018), although the “best” K-value across all populations was higher (see below). The clustering of populations into two genetic groups (K=2) revealed a striking correspondence to geography (Fig. 2C), where the eastern populations (dark blue) are separated from those populations further south and to the west (light blue). This division into two main clades probably suggests some past geographic separation. When three genetic groups (K=3) are considered, the eastern populations are further separated into two clusters, again largely corresponding to geography and reflecting the phylogenetic structure of the data; K=4 distinguishes the west Australia populations from those on the south-eastern coast.

**Fig 2.**
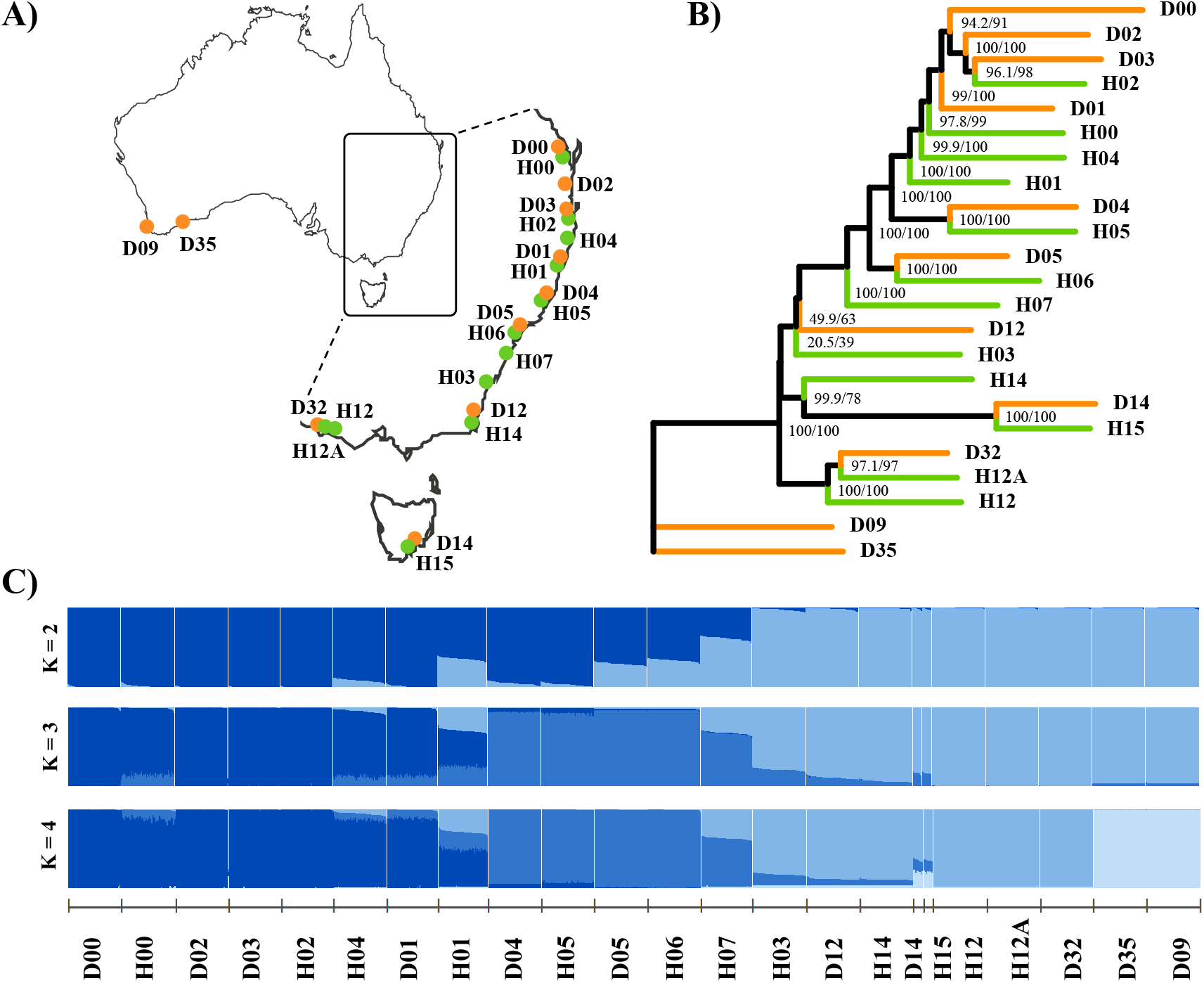
Sampling locations and genetic clustering of *Senecio lautus* populations. **(A)** Sampling locations of the 23 Dune (orange) and Headland (green) *Senecio lautus* populations along the coast of Australia. **(B)** Maximum likelihood phylogeny of Dune and Headland populations implemented in *IQ-TREE*. Numbers on each node represent the SH-alRT support (%), followed by the ultrafast bootstrap support (%). **(C)** Bayesian assignment of individuals to genetic clusters within *fastSTRUCTURE* for K=2-4. Each of the 1,319 individuals is depicted as a bar, with colors representing ancestry proportions to each cluster. Populations are ordered according to their geographic distribution along the coast.

### Minimal admixture across the system

To understand the role of gene flow in shaping the patterns of divergence across the system, we explored patterns of admixture in a phylogenetic context within *TreeMix*. In the absence of migration, the *TreeMix* phylogeny explained 95.9% of the data, with 24 additional migration events augmenting this value to 98.9 % (Supplementary Fig. S1). Fig. 3A shows the first migration event (P < 2.2×10^−308^) with a migration weight (*w*) of 0.40. Although the 24 other migration events were also significant (P_average_ = 2.92×10^−3^, SD = 0.0062), their individual weightings were small (see Supplementary Fig. S2 for 1-10 migration events), most of them were not between parapatric pairs, and the addition of these migration events did not substantially alter the topology from its estimation in the absence of gene flow. Although these results could suggest a potentially complex colonization history including long distance yet rare migration events, these P-values should be treated with caution. This is because model comparisons in *TreeMix* suffer from multiple testing, a large number of parameters, and the estimated graph can be inaccurate (Pickrell and Pritchard 2012). We therefore tested the robustness of this inference using *f3-statistics*. All *f3-statistics* were positive (Supplementary Fig. S3), giving no evidence of admixture between any populations. Strong isolation by distance within each ecotype further supports this contention using 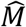 as a proxy for migration rates (IBD within Dunes: Mantel test, r = −0.83, P = 0.0001; within Headlands r = −0.73, P = <0.0001; Fig. 3B). A strong IBD trend also exists between ecotypes for the eight pairs studied here (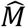: F_1,6_ = 0.55, P = 0.05661, multiple R^2^ = 0.48, Fig. 3B). Although the same trend was seen in the migration rate estimates from *fastsimcoal2*, it was not statistically significant, perhaps due to the low sample size (*fastsimcoal2*: F_1,6_ = 0.66, P = 0.4475, multiple R^2^ = 0.10, Fig. 3C).

**Fig 3.**
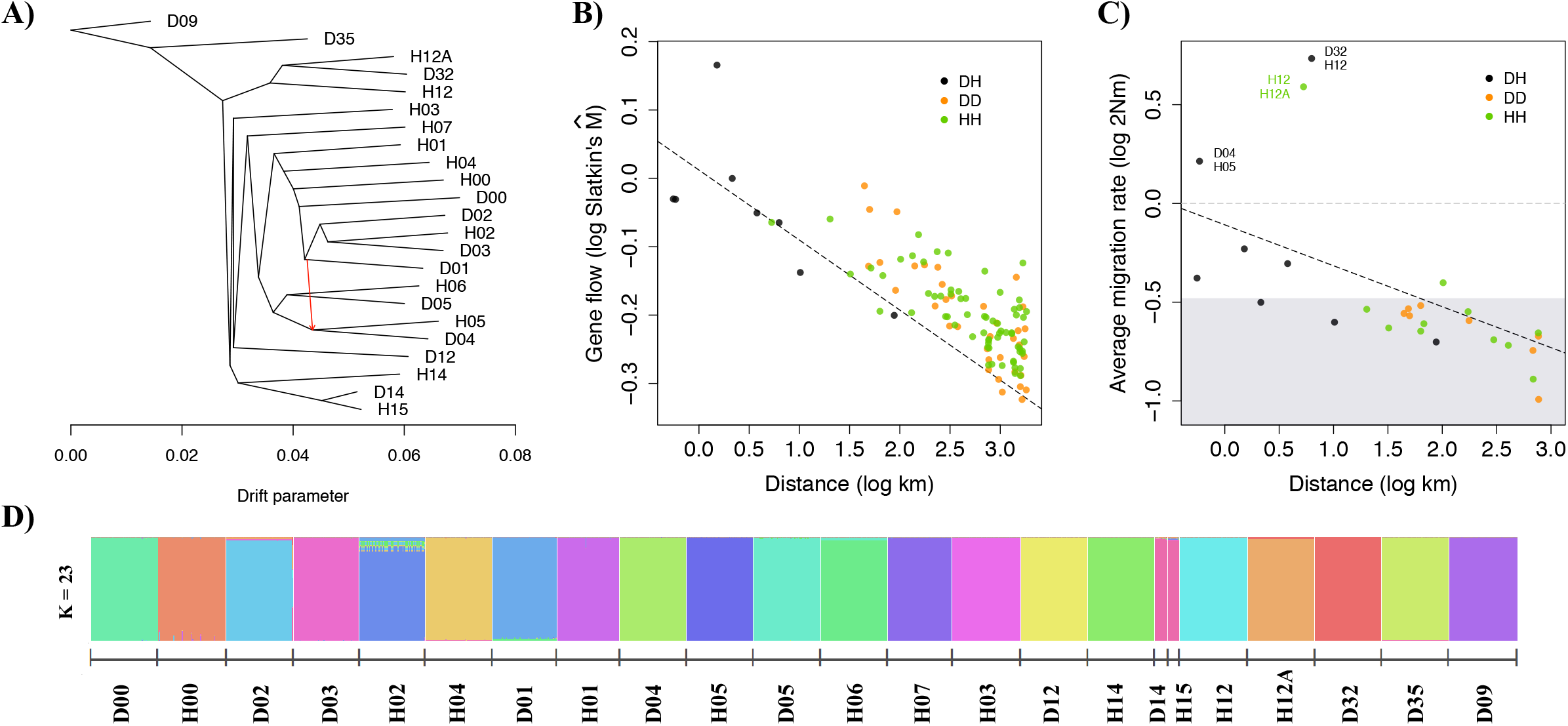
Patterns of long-distance gene flow, isolation by distance, and genetic clustering. **(A)** Maximum likelihood tree with one migration event inferred in *TreeMix*, the x-axis representing genetic drift. The red arrow represents the migration event (*w* = 0.40). **(B)** Patterns of isolation by distance using Slatkin’s 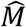 for parapatric Dune-Headland (black), Dune-Dune (orange) and Headland-Headland (green) pairs. Black dashed line represents the linear model for the DH comparisons. **(B)** Patterns of isolation by distance across Dune and Headland populations for Dune-Headland (DH, black), Dune-Dune (DD, orange) and Headland-Headland (HH, green) pairs. Average migration rate is the mean bidirectional migration for each pair estimated in *fastsimcoal2*. Grey shading represents the null expectation for migration rates, inferred from the maximum migration value from three allopatric comparisons (2*Nm* = 0.33). Grey horizontal dashed line represents one migrant per generation (2*Nm* = 1). Pairs falling above this line are labelled. Black dashed line represents the linear model for the DH comparisons. **(D)** Bayesian assignment of individuals to genetic clusters within *fastSTRUCTURE* for K=23. Each of the 1,319 individuals is depicted as a bar, with colors representing ancestry proportions to each cluster. Populations are ordered according to their geographic distribution along the coast.

The absence of admixture across the system is also supported by *fastSTRUCTURE* across all populations. The inferred value of K is close to the number of sampled populations (Supplementary Figs. S4B, S4C) and each population is genetically distinct (Supplementary Fig. S4A), suggesting that *S. lautus* has a simple demographic history with limited admixture (Lawson et al. 2018). Specifically, the K-value that best explained the structure in the data was 22, and the K-value that maximized the marginal likelihood of the data was 28 (Supplementary Fig. S4B). The rate of change in the likelihood of each K-value was negligible for K=24-28 (Supplementary Fig. S4C). Together, this suggests that the optimal K-value is around 23, which is the number of populations within our study. The *fastSTRUCTURE* results for K=23 show that each population forms a distinct genetic cluster (Fig. 3D), suggesting very little, if any, admixture between them. This further implies that each sampled population has been separated from other populations long enough to be genetically distinct (see pairwise F_ST_ values in Supplementary Table S1), and with insufficient levels of gene flow to homogenize their genomes (Lawson et al. 2018). Further, when we examined all K-values from 1-23, we found a distinct hierarchical structure that mirrors the phylogeny, suggesting that such structure is an accurate representation of the history of the populations. The Tasmania population pair (D14-H15) should be treated with caution due to the smaller sample size (n_Dune_ = 12, n_Headland_ = 11) compared to other populations (n_mean_ = 62, SD = 1.19). For groups with few samples, genetic clustering programs such as *fastSTRUCTURE* are likely to assign them as mixtures of multiple populations rather than their own distinct population (Lawson et al. 2018). This is evident for K=22, where the Tasmania populations appear admixed (Supplementary Fig. S4A).

### Minimal gene flow between parapatric ecotypes and distant populations

We investigated whether the parapatric ecotypes at each locality have diverged in the face of gene flow by analyzing patterns of admixture in *STRUCTURE* and directly estimating levels of gene flow in *fastsimcoal2.* We observed very few admixed individuals between the parapatric Dune-Headland populations at each locality within the *STRUCTURE* analysis for K=2 (Fig. 4A). On average, 9.36% (SD = 5.48) of individuals were admixed per population, although their admixture proportions were on average less than 1% (mean = 0.8%, SD = 1.8). This suggests that gene flow between parapatric populations might be rare and ongoing, or might have ceased in the past. For all pairs, the best K-value based on the Evanno method (Evanno et al. 2005) was K=2 (Supplementary Fig. S5). Demographic modelling in *fastsimcoal2* (Fig. 4B) revealed the most likely divergence model for all but three comparisons within and between ecotypes was bidirectional gene flow after secondary contact (mean *w_i_* = 0.99, SD = 0.05; Supplementary Table S4; Supplementary Fig. S6). For comparisons D04-H05, D14-H15 and D04-D05, the most likely divergence model was ancient bidirectional secondary contact (w_i_ = 1.00).

**Fig 4.**
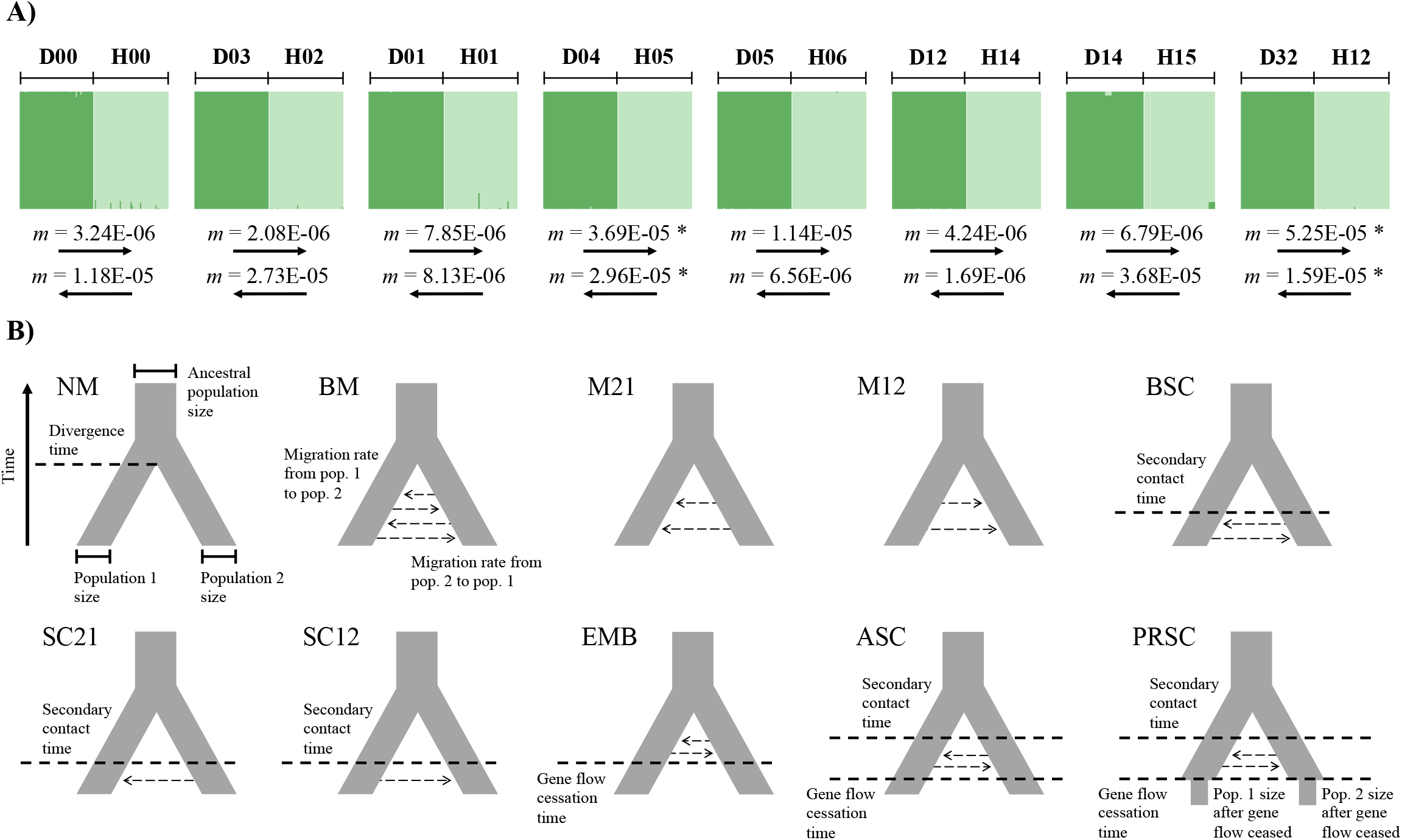
Patterns of gene flow and admixture between parapatric Dune-Headland populations. **(A)** Bayesian assignment of individuals to genetic clusters within *STRUCTURE* for K=2 for the Dune (dark green) and Headland (light green) ecotypes at each locality. Each individual is depicted as a bar, with colors representing ancestry proportions to each cluster. Below are the migration rates (*m*, forward in time) for the most likely divergence model for Dune to Headland, and Headland to Dune within each locality estimated within *fastsimcoal2*. Asterisks denote pairs with 2*Nm* > 1. **(B)** Schematic diagram representing the ten demographic models run in *fastsimcoal2* and their estimated parameters: no migration (NM), bidirectional migration (BM), population 2 to population 1 migration (M21), population 1 to population 2 migration (M12), bidirectional migration after secondary contact (BSC), population 2 to population 1 migration after secondary contact (SC21), population 1 to population 2 migration after secondary contact (SC12), bidirectional migration after population splitting with cessation of gene flow (EBM), ancient bidirectional secondary contact (ASC), population size reduction after bidirectional secondary contact (PRSC).

For Dune-Headland population pairs, direct measurements of migration rates were very low, with all Dune-Headland migration rates below one (2*Nm* < 1.00), except for D04-H05 and D32-H12 where 2*Nm* was above one (Fig. 3C upper section, 4A; Supplementary Tables S2, S3). This is perhaps not surprising given that the populations of D04-H05 are geographically the closest of all parapatric pairs, and the environments of the south-eastern D32-H12 populations are less contrasting than populations in the eastern clade. For Dune-Dune comparisons we also detected very low migration rates (2*Nm*_mean_ = 0.24, SD = 0.09; Supplementary Tables S2, S3), with all comparisons containing 2*Nm* < 1.00. Similarly, for Headland-Headland comparisons we again detected very low migration rates (2*Nm*_mean_ = 0.57, SD = 1.08), with all comparisons containing 2*Nm* < 1.00, with the exception of H12-H12A (Fig. 3C upper section; Supplementary Tables S2, S3). Across all comparisons, all Dune-Dune pairs and most Headland-Headland pairs exhibited gene flow levels lower than the maximum migration rate of allopatric populations separated by more than 1,500 km (i.e., the null gene flow expectation; 2*Nm* = 0.33; Fig. 3C). Three Dune-Headland pairs (D00-H00, D03-H02 and D12-H14) were also within this null range. Alternative models that assumed very low gene flow, while keeping other demographic parameters fixed, were not a better fit for the data with the exception of the D03-H02 pair (Supplementary Fig. S7). Although the most likely divergence scenario for all population comparisons was bidirectional gene flow after secondary contact and ancient bidirectional secondary contact, migration rates under all models were very low across all population pairs (2*Nm*_mean_ = 0.68, SD = 1.52, Supplementary Table S4). Thus, even if our choice of model was biased towards secondary contact or models with migration (Roux et al. 2016; Momigliano et al. 2021), the extent of gene flow during the history of populations is consistently low and does not strongly depend upon the mode of divergence.

### Potential for gene flow to obscure a single origin scenario

We analyzed a neutral coalescent model representing a single origin of the derived ecotype to investigate under which situations the history at a neutral locus unlinked to the selected site would indicate a pattern of non-monophyly of the derived ecotype, thus potentially supporting a false inference of parallel evolution. We found a clear influence of all examined parameters (Fig. 5A) on the probability of inferring non-monophyly, suggesting that certain demographic scenarios can lead to an observed phylogenetic signal that belies the history of a single origin of ecotypes. When internal branches (*t2* – *t1*) are short, incomplete lineage sorting will cause higher probability of non-monophyly, and this is not much increased by migration (Fig. 5B, lower section of graph). When internal branches are long, the probability of non-monophyly is low, and only somewhat more sensitive to increasing migration rate (Fig. 5B, upper section of graph). Long terminal branches (*t1*) also influence the probability of non-monophyly; the more diverged populations contributing migrants are from one another prior to migration into the other ecotype, the more this increases the chances of non-monophyly (Fig. 5C, upper section of graph). This reflects the fact that the introgressed alleles in different recipient populations will have experienced more independent drift prior to the introgression event. When the terminal branches are short relative to the timing of migration, very high levels of migration are required to have a high probability non-monophyly (Fig. 5C, lower section of graph). Note that the probability of non-monophyly is more sensitive to the length of the internal branches, as these determine the likelihood of incomplete lineage sorting (see the different scale of probability in Fig. 5C compared to Fig. 5B). Simulation of a single origin scenario with continuous gene flow in *SLiM2* (Haller and Messer 2017) showed patterns consistent with the above coalescent results (Supplementary Fig. S8A).

**Fig 5.**
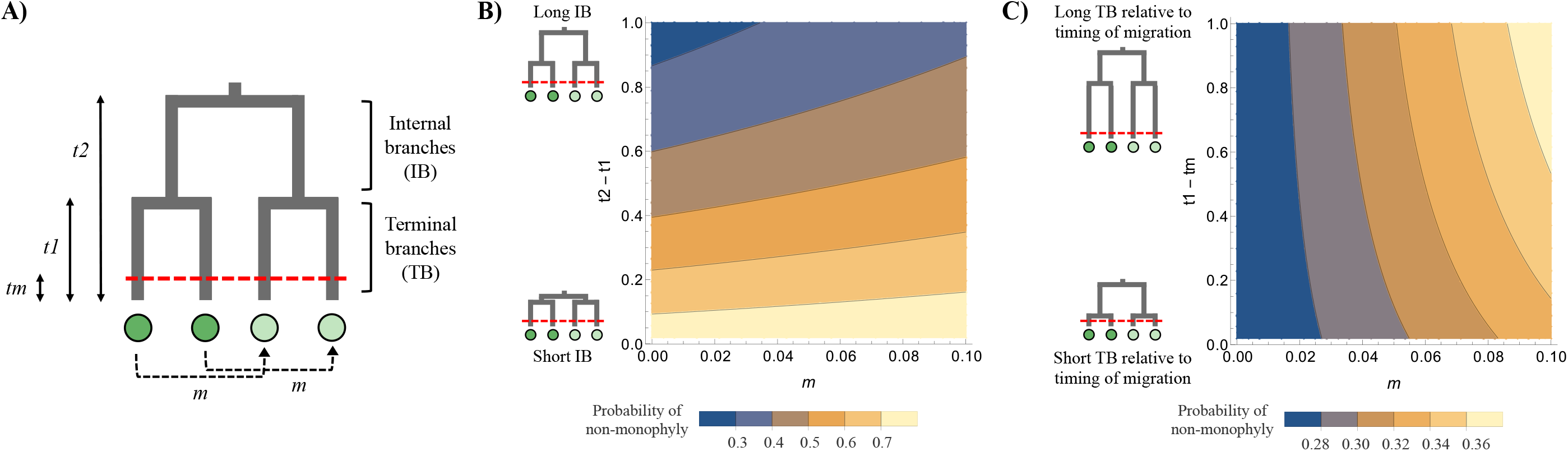
Coalescent modelling to infer the probability of non-monophyly. **(A)** Schematic diagram representing the modelled single origin scenario. *t2* represents the time to the split of the ancestral population (i.e., the initial origin of the Dune and Headland ecotypes). Each of these two ecotypes further split at time *t1* in the past. Parapatric populations at each locality are connected with dashed lines, and an instantaneous burst of migration (*m*) occurs at time *tm* in the past (dashed horizontal red line). In the model, all times are expressed in units of 2*N* generations. Light green and dark green circles represent populations from different ecotypes. **(B, C)** Probability of non-monophyly, falsely suggesting the parallel evolution of ecotypes. *tm* is set 0.1 (corresponding to 50,000 generations for a diploid population size of 250,000). Small phylogenies are schematic diagrams of the extreme values of the y-axis. **(B)** High probability of non-monophyly occurs when internal branches (IB) are short (lower yellow region). *t1* is set to 1 (corresponding to 500,000 generations for N=250,000). **(C)** The probability of non-monophyly requires high migration and increases when length of terminal branches (TB) are longer prior to the burst of migration (right-hand yellow region). *t2* is set to 1 (corresponding to 500,000 generations). Note the different scale of probability in panel **(C)** compared to panel **(B)**.

Although we cannot directly map our observed data for *S. lautus* in the modelled or simulated parameter spaces, we can nonetheless explore potential evidence of phylogenetic distortion by examining whether the divergence time estimates from *fastsimcoal2* (which accounts for gene flow) are in accordance with the phylogenetic topology estimated in *IQ-TREE* (which does not account for gene flow). This was the case for our data: we observed deeper divergence times for populations of the same ecotype compared to sister-taxa of different ecotypes. For D04-H05 and D05-H06, the average divergence time between populations of the same ecotype (D04-D05 and H05-H06) was 83,846 generations (SD = 3,021), whereas the average divergence time between populations at each locality (D04-H05 and D05-H06) was 52,569 generations (SD = 21,719). This was also true for D14-H15 and D32-H12, where the average divergence time between populations of the same ecotype (D14-D32 and H12-H15) was 68,723 generations (SD = 17,526), and the average divergence time between populations at each locality (D14-H15 and D32-H12) was 49,882 generations (SD = 15,805). Finally, we would expect that had gene flow been substantial we would observe high F_ST_ values between parapatric populations, but our estimates are near 0.2.

## Discussion

We have used an array of complementary approaches to disentangle the demographic history of the coastal *Senecio lautus* ecotypes. In this system, many lines of evidence support a multiple origin scenario for the evolution of the parapatric Dune and Headland populations. The demographic history of this system reveals striking population structure and a strong effect of geography and restricted dispersal, to the extent that all populations are evolving largely independently from each other. Together with previous results from transplant experiments (Melo et al. 2014; Richards et al. 2016; Richards and Ortiz-Barrientos 2016; Walter et al. 2016; Walter, Wilkinson, et al. 2018; Wilkinson et al. 2021), our results convincingly show that selection and drift, rather than gene flow, play a predominant role in the genetic structure among ecotypic populations in this system. Below we discuss these results considering parallel parapatric divergence in this highly replicated system.

### Strong genetic structure between ***Senecio lautus*** coastal populations

Large scale genetic structure of *S. lautus* clusters populations according to their geographic distribution along the Australian coast, and not by the environment they occupy. Within *fastSTRUCTURE*, the largest genetic groups encompass two clades which are largely independent of each other, do not show evidence of long-distance gene flow between them, and also appear to contain multiple repeated instances of parapatric divergence. This genetic structure, where populations group by geography and not ecology, is mirrored in the phylogeny, and is consistent with our previous work using targeted sequencing of 13 neutral genes (Melo et al. 2019) and RADseq using pools of individuals (Roda, Ambrose, et al. 2013). There is also a strong signal of isolation by distance (Wright 1943) within each ecotype as well as across Dune-Headland pairs, implying long distance dispersal within the system is not pervasive, which might indicate that populations are at an equilibrium between dispersal and drift (Slatkin 1993). This is perhaps not surprising given that Dune and Headland populations have restricted geographic ranges along the coast.

Fine scale genetic structure at the level of the locality (i.e., parapatric Dune-Headland populations) shows that each population is genetically distinct, with little admixture, as suggested by *fastSTRUCTURE* and *f3-statistics*. Despite the high potential for gene flow between parapatric populations due to their close geographic proximity and relatively weak F1 intrinsic reproductive isolation (Melo et al. 2014; Richards et al. 2016; Walter et al. 2016), multiple transplant experiments in the system have shown that divergent natural selection is strong and creates extrinsic reproductive isolation between Dune and Headland populations at each locality (Melo et al. 2014; Richards et al. 2016; Richards and Ortiz-Barrientos 2016; Walter et al. 2016; Walter, Wilkinson, et al. 2018; Wilkinson et al. 2021). Therefore our findings are in agreement with theoretical expectations, where parapatric divergence and speciation is favored when gene flow is limited and selection against immigrants and hybrids is strong (Bank et al. 2012; Blanckaert and Hermisson 2018). Overall, a combination of strong selection and limited dispersal can explain why parapatric populations persist despite the opportunity for homogenizing gene flow between them.

### Parallel evolution of parapatric ***S. lautus*** ecotypes with minimal levels of gene flow

A common doubt arising in purported cases of parallel evolution is whether gene flow is responsible for the grouping of populations by geography and not by ecology (Quesada et al. 2007; Johannesson et al. 2010; Bierne et al. 2013; Butlin et al. 2014; Martin et al. 2015; Rougemont et al. 2015; Le Moan et al. 2016; Meier et al. 2017; Pérez-Pereira et al. 2017; Trucchi et al. 2017; Rougeux et al. 2019). A single origin of ecotypes combined with high levels of gene flow between parapatric ecotypes at each locality can alter the phylogenetic relationships of populations, falsely suggesting multiple independent origins. This is because genetic structure at neutral markers can be decoupled from colonization history via introgression and incomplete lineage sorting (Endler 1977; Barton and Hewitt 1985; Coyne and Orr 2004; Bierne et al. 2013). This needs careful scrutiny in our system as previous work showed that genomic divergence was more heterogenous in parapatric populations compared to allopatric populations (Roda, Ambrose, et al. 2013), a signature of divergence with gene flow (Feder and Nosil 2010; Feder et al. 2012). However, this pattern can also arise due to ancestral polymorphism and incomplete lineage sorting if parapatric populations are younger than allopatric populations (Cruickshank and Hahn 2014), which our current work shows. Thus, processes unrelated to divergence in the face of high levels of gene flow could have also contributed to the patterns of genomic divergence in this system (Roda, Ambrose, et al. 2013).

To help understand the role of gene flow during parapatric divergence in *S. lautus*, we first directly estimated rates of gene flow within *fastsimcoal2*. Unexpectedly, we observed minimal levels of gene flow within parapatric *S. lautus* Dune-Headland pairs. This reveals that previous patterns of genomic divergence among these populations (Roda, Ambrose, et al. 2013) likely reflect a signature of increased genome-wide differentiation over time in allopatry, and incomplete lineage sorting in parapatric populations rather than heterogeneous divergence in the face of high levels of gene flow. Furthermore, we observed that most Dune-Headland levels of bidirectional gene flow were similar to populations from different clades and separated by greater than 1,500km, suggesting that most Dune and Headland populations at each locality could be viewed as effectively allopatric. Although unmodelled changes in population size tend to favor secondary contact models of gene flow (Momigliano et al. 2021) and our assumed genomic homogeneity of migration might affect the relative ranking of demographic models (Roux et al. 2016), our estimated migration rates were consistently very low across most models of gene flow (see Supplementary Table S4 for details). We do not place emphasis on distinguishing between the different models of migration, but rather whether parapatric divergence occurred in the presence or absence of gene flow – we are confident that a small amount of gene flow has occurred between the ecotypes at most localities. There are two notable exceptions (D04-H05 and D32-H12), which experience levels of gene flow that would make them genetically indistinguishable (2*Nm* > 1.00) (Slatkin 1985a). Thus, it is still possible that gene flow has altered the observed phylogenetic topology for these populations.

We therefore explored the conditions which are likely to obscure the history of colonization by modelling the neutral divergence process through coalescent analyses and simulations in *SLiM2*. We observed that the likelihood of non-monophyly is accentuated with very short internal branches, which will lead to high levels of retained ancestral polymorphism and therefore increase the probability that true sister taxa do not remain monophyletic; this effect is largely independent of migration rates as gene flow will not contribute more to population similarity beyond what ancestral polymorphism already does. Short internal branches are frequently detected in systems where diversification occurs rapidly, such as in cases of adaptive radiations (Schluter 2000), which seems to be the case in *S. lautus*. Our theoretical approach also revealed that increasing levels of gene flow increases the likelihood of non-monophyly, especially when the time of migration is further from the second population split. This is because alleles introgressing into different populations will have experienced more independent drift. More importantly, in these cases of secondary contact, quite high levels of migration are required to create the appearance of multiple origins. Although conventional thinking highlights that even small amounts of gene flow have the potential to mix populations and erode their history (e.g., Endler 1977; Barton and Hewitt 1985; Coyne and Orr 2004), our work suggests that this picture can be more complex in cases of secondary contact involving multiple population pairs. As expected, our work reveals that it is important to consider the joint contributions of gene flow as well as ancestral polymorphism when inferring the likelihood of phylogenetic distortion.

Within *S. lautus*, even though the short internal branches and recent secondary contact have the potential to obscure the phylogeny and falsely suggest parallel evolution, this is likely circumvented by the extremely low rates of gene flow between most parapatric ecotypes. In other words, it appears that higher amounts of gene flow would be needed to counteract the divergence that has accumulated over time in the *S. lautus* system. We must also note that our theoretical approach is conservative as we have ignored the effects of selection against introgressed alleles. We expect that linkage to loci underlying local adaptation should act to decrease the probability of a phylogeny topology switch at the locus considered. As such, a polygenic basis of local adaptation could greatly reduce the probability of a topology switch due to gene flow. When considering our theoretical work in combination with patterns of gene flow and genetic structure in the system, there is strong evidence to suggest that *S. lautus* populations have evolved multiple times independently.

Further evidence that gene flow has not obscured a single origin scenario in *S. lautus* comes from comparing joint estimates of gene flow and divergence times (as implemented in isolation with migration models) between population pairs of the same ecotype and putative sister populations of divergent ecotypes. We observed that populations of the same ecotype consistently show deeper divergence times than those from different ecotypes, which reflects the topology of the phylogeny estimated in the absence of gene flow. In addition, constructing the phylogeny considering gene flow (in *TreeMix*) did not alter the topology from its estimation in the absence of gene flow. Although the divergences at two localities (D04-H05 and D32-H12) experience levels of gene flow high enough to potentially result in phylogenetic distortion, their levels of differentiation are rather high. Furthermore, each of these pairs is from a separate clade and are genetically isolated from other such pairs, so even moderate levels of gene flow within each pair would not have distorted the phylogeny across the entire system. Together, these results imply that phylogenetic distortion is highly unlikely in *S. lautus* and that such relationships reflect the true history of populations and ecotypes.

Additional support for the parallel evolution of *S. lautus* populations comes when considering our results in combination with previous work. In *S. lautus* a clear association between environment and phenotype exists in both common garden (Walter, Aguirre, et al. 2018) and field conditions (James et al. 2020), as well as strong divergent selection within each population pair (Melo et al. 2014; Richards et al. 2016; Richards and Ortiz-Barrientos 2016; Walter et al. 2016; Walter, Wilkinson, et al. 2018; Wilkinson et al. 2021). This striking association between trait and environment implies that selection would have had to independently resist the introduction of maladaptive alleles to repeatedly regenerate the optimal phenotype across multiple populations. Thus, parapatric systems where gene flow may be extensive can still be considered ‘parallel’ because natural selection independently maintains differentiation between ecotypes at each locality. In this manner, Lee and Coop’s (2019) framing of independence as an overlap in selective deaths across populations can be extended to consider both the overlap in selective deaths during the initial sweep, as well as during a secondary phase of resisting maladaptive gene flow. Within *S. lautus*, the current and previous work provides strong support not only for the parallel origin of ecotypes, but the parallel action of natural selection. Furthermore, similar phenotypes across *S. lautus* replicate populations have arisen via mutations in different genes (Roda, Liu, et al. 2013; James et al. 2020; Wilkinson et al. 2021), providing even further evidence that natural selection would have acted independently within each population. Overall, our results indicate that *S. lautus* is a highly replicated system of parapatric divergences.

### The limits of inference from parallel evolution

The *S. lautus* system allows us to study the deterministic nature of parallel evolution in multiple ways. For instance, we can now begin to understand how genetic architectures vary and evolve during adaptation. In doing so, we can then demonstrate whether alleles, genes or pathways have been repeatedly selected for across replicate populations (Lee and Coop 2017; Lee and Coop 2019). This will also help us understand whether adaptation arises from new mutations or standing genetic variation (e.g., Lee and Coop 2017), or from fixation of functionally equivalent alleles (such as during polygenic adaptation; Berg and Coop 2014; Tiffin and Ross-Ibarra 2014; Yeaman 2015), or loss-of-function mutations (e.g., Chan et al. 2010). Once adaptive genes have been identified, studies of parallel evolution should directly link the adaptive loci to phenotypic traits and further demonstrate that the traits themselves have been under repeated selection in independent populations (Storz and Wheat 2010; Pardo-Diaz et al. 2015; Hoban et al. 2016). In systems where this method is not feasible, our study demonstrates that studying genome-wide loci can still uncover patterns of phylogeography and migration that are consistent with parallel evolution.

Finally, in our work we have unusually high power to detect gene flow, as the number of individuals sequenced in each of our populations is large (*N*_mean_ = 57, 2*N*_mean_ > 100 chromosomes per population). This sampling regime allowed us to sample many rare variants and therefore better distinguish ancestral polymorphism from migration. Studies undertaking demographic modelling often sample 10-25 individuals per population (e.g., Roesti et al. 2015; Kautt et al. 2016; Trucchi et al. 2017) and occasionally even less than 10 (e.g., Meier et al. 2017). Thus these studies cannot easily distinguish shared variants due to gene flow from ancestral polymorphism, which can make results biased to detecting moderate to high levels of gene flow, especially for recently diverged populations and in underpowered studies (Slatkin 1985b; Hey and Nielsen 2007; Strasburg and Rieseberg 2010; Cruickshank and Hahn 2014). As our coalescent modelling revealed that only a small amount of gene flow is needed to obscure a phylogenetic topology under certain conditions (e.g., during very young adaptive radiations), studies that estimate levels of gene flow with few numbers of individuals and loci should treat results with caution.

Overall, we provide strong evidence for multiple origins of parapatric Dune and Headland populations within *Senecio lautus*. Across this highly replicated system we observed phylogenetic clustering by geography, strong genetic structure between populations, isolation by distance, and surprisingly low gene flow between parapatric populations at each locality as well as in the system as a whole. Coalescent modelling and simulations confirmed that levels of gene flow are likely not high enough to obscure a single origin scenario in *S. lautus*. Furthermore, the phylogenetic relationships of populations estimated in the presence of gene flow agree with the main phylogeny, which supports a multiple origin scenario. These results from our current work in combination with strong divergent selection between ecotypes (Melo et al. 2014; Richards et al. 2016; Richards and Ortiz-Barrientos 2016; Walter et al. 2016; Walter, Wilkinson, et al. 2018; Wilkinson et al. 2021), strong trait-environment association in the system (Walter, Aguirre, et al. 2018; James et al. 2020) and adaptation across replicate populations occurring mainly via mutations in different genes (Roda, Liu, et al. 2013; James et al. 2020; Wilkinson et al. 2021), implies that selection has independently driven the parallel evolution of populations. This makes *S. lautus* one of the clearest examples of the parallel evolution of ecotypes discovered yet, adding to the increasing number of potential cases of parallel evolution in plants (Foster et al. 2007; Trucchi et al. 2017; Cai et al. 2019; Konečná et al. 2019). It also positions the species as a powerful system of replicated parapatric divergence to study the origin of adaptation and reproductive isolation.

## Methods

### Sample collection and DNA extraction

Leaf samples for DNA extraction were collected from 23 Dune and Headland populations of *Senecio lautus* along the coast of Australia, which included eight parapatric Dune-Headland population pairs, three allopatric Headland populations, and three allopatric Dune populations (n_mean_ = 58, n_total_ = 1338; Fig. 2A, Supplementary Table S5). We sampled mature (flowering) plants evenly across the geographic range of each population, ensuring that sampled plants were at least one meter apart. Leaf collection was undertaken with permission from the Redland City Council, New South Wales National Parks and Wildlife Services (Permit number: SL100215), Victoria Department of Environment and Primary Industries (Permit number: 10007220), Tasmania Department of Primary Industries, Parks, Water and Environment, and the Western Australia Department of Parks and Wildlife. DNA was extracted using a modified CTAB protocol (Clarke 2009) and cleaned with Epoch Life Sciences spin columns. We quantified sample concentration with the Invitrogen Quant-iT PicoGreen dsDNA Assay Kit, and used the BioTek Take3 Micro-Volume Plate to ensure DNA samples were pure. Samples were standardized to 10ng/μL.

### GBS library construction

We created reduced representation libraries to obtain restriction site associated DNA (RAD) markers. Specifically, we used a two-enzyme Genotyping-by-Sequencing (GBS) approach (modified from Poland et al. 2012). We created seven libraries from the 23 Dune and Headland populations, each containing 192 barcoded individuals. For each individual, genomic DNA was digested with the restriction enzymes Pst1-HF (New England Biosciences; NEB) and Msp1 (NEB). Forward and reverse barcodes were ligated to fragments from each sample, and subsequently cleaned with homemade Serapure beads (Faircloth and Glenn 2011; Rohland and Reich 2012). For each sample we amplified the fragments and added Illumina sequencing primers via PCRs with 14 cycles. Each sample was quantified with the Invitrogen Quant-iT PicoGreen dsDNA Assay Kit. We created seven equimolar pools (192 individuals per pool), ensuring each population was evenly distributed across the pools. Each pool was size-selected using the BluePippin (2% DF Marker V1, 300-500bp; Sage Science), and cleaned with the Monarch PCR & DNA cleanup kit (NEB). Pooled libraries were sent to the Beijing Genomics Institute (BGI) for sequencing on seven lanes of the HiSeq4000, using 100bp paired-end sequencing.

### Reference genome

A PacBio *S. lautus* reference genome (v1.0) was created from a selfing Headland individual from Lennox Head (see Wilkinson et al. (2021) for additional details of the individual). The plant was put in darkness for three days before young leaves were snap frozen in liquid nitrogen and sent to BGI for DNA extraction and PacBio long-read sequencing (56 SMRT cells). A total of 11,481,638 filtered subreads (5,072,756 from RSII sequencer and 6,408,882 from Sequel sequencer) were used with a mean read length of ~8K base pairs and an expected genome coverage of ~60X. The genome was de novo assembled using CANU 1.6 (REF) with default settings. The final assembly consisted of 60,947 contigs with an N50 of 34K. The BUSCO gene content completeness was 86% (4% fragmented and 10% missing).

### Bioinformatics

For the GBS sequencing, BGI removed forward barcodes and quality filtered the raw reads to remove reads containing Illumina adaptors, low quality reads (> 50% of bases < Q10), and reads with > 10% Ns. We trimmed reverse barcodes with *TagCleaner* standalone v0.12 (Schmieder et al. 2010). We retained an average of 2,849,159 clean reads (SD = 827,036) across the 1,319 individuals (after the removal of 19 individuals with high levels of missing data, see below; Supplementary Table S6). Reads were mapped to the *S. lautus* reference PacBio genome v1.0 with *BWA-MEM* v0.7.15 (Li and Durbin 2009; Li 2013). On average, 86% of reads (SD = 15) mapped to the reference genome, and 81% (SD = 15) mapped properly with their paired-read (Supplementary Table S6). *PicardTools* v2.7.0 (Broad Institute 2019) was used to clean aligned reads and to add read groups (PCR duplicates were not marked for removal). We jointly called all variant and invariant sites for each population with *FreeBayes* v1.1.0 (Garrison and Marth 2012). Because SNPs were separately called for each of the 23 populations, we first normalized the 23 VCF files before merging them together. This was achieved by first using BCFtools v1.4.1 (Li et al. 2009) to split multiallelic sites into biallelic records. Each file was then normalized by re-joining biallelic sites into multiallelic records. We then left-aligned and normalized indels, and used *vt* (Tan et al. 2015) to decompose biallelic block substitutions into separate SNPs for each population. We merged the 23 per-population VCF files into one large file for subsequent SNP filtering.

We largely followed the *dDocent* pipeline for SNP filtering (Puritz, Hollenbeck, et al. 2014; Puritz, Matz, et al. 2014), including iterative filtering to maximize the number of sampled SNPs (O'Leary et al. 2018). Using *VCFtools* v0.1.15 (Danecek et al. 2011), we first retained sites if they were present in > 50% of individuals, had a minimum quality score of 30, and a minimum minor allele count of 1. We filtered for a minimum depth of 3 for a genotype call. Individuals were removed if they contained > 40% missing data. We then filtered for a maximum mean depth of 100, and a minimum mean depth of 10. We filtered for missing data per population, removing sites if they contained > 50% of missing data within each population. We refiltered for an overall missing data of 20%. Indels were removed with *vcflib* (Garrison 2016). We then filtered for population-specific Hardy Weinberg Equilibrium using the *filter_hwe_by_pop.pl* script within *dDocent*. See below for the minor allele frequency thresholds for each analysis.

### Do populations cluster by geography or ecotype?

To explore the broad patterns of genetic clustering of populations, we performed two separate analyses: phylogeny construction and *fastSTRUCTURE* (Raj et al. 2014). We used *PLINK* v1.9 (Purcell et al. 2007) to remove SNPs with a minor allele frequency (MAF) of < 0.05, and also to thin SNPs by retaining one unlinked SNP per RAD locus. This dataset contained 3,844 unlinked SNPs across the 1,319 individuals. We generated a maximum likelihood phylogeny within *IQ-TREE* v1.6.0 (Nguyen et al. 2015) using the polymorphisms-aware phylogenetic model (Schrempf et al. 2016). We first used ModelFinder (Kalyaanamoorthy et al. 2017) to determine the best-fit substitution model for the data (TVMe+FQ+P+N9+G4), and increased the virtual population size (N) to the maximum value of 19 (as recommended by Schrempf et al. 2016). Default parameters were used for tree construction, with the western Australia D09 population assigned as the outgroup. To assess convergence, we undertook 10 separate runs of *IQ-TREE* and examined tree topology (which remained unchanged across the 10 independent runs). We also ensured that the log-likelihood values were stable at the end of each run. Branch support was performed using 10,000 replicates of UFboot (Hoang et al. 2018), and 10,000 replicates of SH-aLRT (Guindon et al. 2010).

We further explored broad patterns of population structure using the variational Bayesian framework, *fastSTRUCTURE* v1.0 (Raj et al. 2014). Here, we implemented *fastSTRUCTURE* to provide extra evidence for whether populations genetically cluster by geography or ecotype. We did not infer specific historical admixture scenarios from *fastSTRUCTURE*, as different demographic scenarios can give rise to indistinguishable structure plots (Lawson et al. 2018). The *fastSTRUCTURE* algorithm assigns individuals into genetic clusters (K) by minimizing departures from Hardy-Weinberg equilibrium and inferring individual ancestry proportions to each genetic cluster. We followed the recommendations by Gilbert et al. (2012) and Janes et al. (2017) and ran the simple prior (K=1-30) with 100 independent runs per K-value. In order to determine the most likely number of genetic clusters (the optimal K), we used the *chooseK.py* script from *fastSTRUCTURE* to examine (1) the K-value that best explained the structure in the data (the smallest number of model components that accounted for almost all of the ancestry in the sample), and (2) the K-value that maximized the marginal likelihood of the data. Results were summarized and plotted in the R package *pophelper* v2.2.7 (Francis 2017).

### Has gene flow shaped patterns of divergence across the system?

To explore patterns of gene flow in a phylogenetic context, we used *TreeMix* v1.13 (Pickrell and Pritchard 2012). *TreeMix* constructs a bifurcating maximum likelihood tree, identifies populations that are poor fits to the model, and sequentially adds migration events that improve the fit of the data. SNPs for which MAF < 0.01 were filtered, which retained 24,933 SNPs across the 1,319 individuals. We constructed an initial 25 maximum likelihood trees with no migration, 1,000 bootstrap replicates in blocks of 50 SNPs with D09 as the assigned outgroup, and selected the tree with the highest log-likelihood as the input tree for all subsequent analyses. We then tested between 1-25 migration events in blocks of 50 SNPs. Trees and migration events were robust to varying the size of the linkage blocks as well as the MAF threshold of the dataset (data not shown). To select the number of migration events, we examined the log-likelihoods and cumulative variance explained by each model, and performed jackknife estimates to obtain the standard error and significance of the weight of each migration event. However, the interpretation of these P-values should be treated with caution due to possible errors in the tree structure as well as the inference of incorrect migration events (Pickrell and Pritchard 2012).

To more formally test for genetic admixture, we used the *threepop* function in *TreeMix* to calculate *f3*-statistics (Reich et al. 2009). The *f3*-statistic determines whether a particular population (*A*) is the result of admixture between two other populations (*B* and *C*). It measures the difference in allele frequencies between populations *A* and *B*, and populations *A* and *C*. Only when admixture is present, do we expect the allele frequency of population *A* to be intermediate between the allele frequencies of populations *B* and *C*. In contrast, in the absence of gene flow, the allele frequency of population *A* should not be consistently intermediate between *B* and *C*. Therefore, *f3* can be interpreted as the amount of shared genetic drift between two populations from a common ancestor. In the absence of admixture, *f3* (*A*; *B*, *C*) will be positive, whereas a significantly negative value of *f3* provides evidence for *A* being admixed from *B* and *C*. We calculated *f3* for all possible triads of populations with jackknifing in blocks of 50 SNPs to obtain Z-scores for calculating statistical significance (Z-score < −3.8 = P < 0.0001).

The erect phenotype is common across Australian species of the genus *Senecio* (Thompson 2005), except for the prostrate *S. lautus* Headland ecotype and a few Alpine populations, suggesting these prostrate forms are derived. We tested for isolation by distance (IBD; Wright 1943) in the ancestral and derived ecotypes to evaluate similarities in their dispersal dynamics (Slatkin 1993). We tested for IBD using migration rates (2*Nm*) inferred in *fastsimcoal2* (see below) as well as Slatkin’s 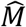, *(1 / F _ST_ − 1)/4*, as a proxy for gene flow (Slatkin 1993). For Slatkin’s 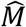, we excluded the Western Australia populations (D09 and D35), and SNPs for which MAF < 0.05 were filtered. We calculated pairwise F_ST_ in VCFtools. We calculated pairwise geographic distances using the following formula, which uses the spherical law of cosines to consider the curvature of the earth:

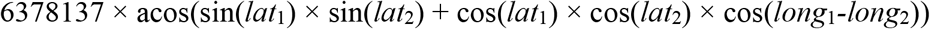

where: 6378137 is earth’s radius in meters, and *lat* and *long* are the latitude and longitude (in radians) of the two populations being compared. For the *fastsimcoal2* migration rates, we tested for IBD between the Dune and Headland within each population pair using a linear model in R v3.4.2 (R Core Team 2017), using an average of the bidirectional gene flow rates for each pair (log-log scale). For Slatkin’s 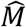 we also tested for IBD between the Dune and Headland of each population pair (log-log scale) using a linear model in R, and tested for IBD within the Dunes, and within the Headlands (log-log scale) using Mantel tests with 9,999 permutations in R (*mantel* function in the *vegan* package; Blanchet et al. 2018).

### Is there gene flow between parapatric populations?

We examined levels of admixture between parapatric populations with *STRUCTURE* v2.3.4 (Pickrell and Pritchard 2012). *STRUCTURE* is a Bayesian MCMC approach that assigns populations into genetic clusters (K) based on individual genotypes by assuming Hardy-Weinberg Equilibrium within a population. It assigns each individual an admixture coefficient to depict the proportion of the genome that originated from a particular K cluster. To increase the numbers of SNPs, we took a subset of the data by excluding the two populations from the west coast of Australia (D09 and D35). Excluding these most divergent populations decreased the amount of missing data and thus increased the number of common SNPs in the south-eastern populations. We used the same filtering procedure as above, removed SNPs with MAF < 0.05 and thinned SNPs in *PLINK* to retain one SNP per RAD locus. Each Dune-Headland population pair was extracted, and SNPs with MAF < 0.05 were filtered. We retained between 837 and 2,606 unlinked SNPs per pair (mean = 1,905 SNPs; SD = 575). *STRUCTURE* analysis was run using the admixture model and the correlated allele frequency model (Falush et al. 2003) with 10 independent runs for K=1-6 (50,000 burn-in and 200,000 MCMC iterations). We visually inspected summary statistics of MCMC runs to ensure convergence of model parameters. As we were specifically interested in detecting admixed individuals between the two ecotypes, we plotted results for K=2. To explore any additional genetic structure within a pair, we also estimated the optimal K-value with the Evanno method (Evanno et al. 2005), by examining the maximum value for ΔK (the second order rate of change in the log probability of data between successive K-values). The R package *pophelper* was used to calculate the ΔK, summarize results and plot the data.

We directly estimated levels of gene flow between population pairs from the site frequency spectrum (SFS) using the composite-likelihood method implemented in *fastsimcoal2* v2.6.0.3 (Excoffier et al. 2013). The joint SFS of two populations is sensitive to demographic processes. For instance, gene flow will result in more low-frequency shared polymorphisms than expected under a non-migration scenario (Hahn 2018). We tested ten demographic models (Fig. 4B), and inferred migration rates, as well as other demographic parameters including current and past population sizes, ancestral population size, divergence time, time of secondary contact, and gene flow cessation time, for eight Dune-Headland (DH) population pairs. We additionally asked whether gene flow was occurring in a linear fashion down the coast within each ecotype by testing eight Dune-Dune (DD) and eleven Headland-Headland (HH) pairs. To determine the baseline level of gene flow inferred by *fastsimcoal2* between isolated populations, namely the null gene flow expectation, we estimated migration rates for three very divergent allopatric populations (>1,500km apart, between the eastern and south-eastern clades; D03-D32, D03-H12, and H02-H12), and took the highest detected migration rate from these allopatric comparisons as the baseline migration rate.

As above, the Western Australia populations (D09 and D35) were excluded from this dataset to increase the number of sampled SNPs. For each pair, we filtered for a minor allele count of one (MAC1), retaining between 6,679 and 19,951 variable sites per pair (mean = 12,155 SNPs, SD = 3,316). By using a MAC1 and a relatively high number of samples per population (mean = 57, SD = 15), we retained rare alleles that are informative about migration events between the populations (Slatkin 1985b). We note that we also ran *fastimcoal2* removing SNPs with MAF < 0.05. We found that migration rates were higher using MAC1 compared to MAF0.05 which means that if anything, we would be overestimating migration rates, which would make our case for very low gene flow even stronger (result not shown). Since we cannot distinguish ancestral from derived alleles, we used the minor allele SFS (folded SFS). We used an *ad hoc* approach to estimate the number of monomorphic sites (Supplementary Methods S1). Gene flow estimates were robust to varying the number of monomorphic sites (data not shown). We generated the joint folded SFS per population pair without downsampling using *vcf2dadi* and *dadi2fsc.2pop* R functions from Liu et al. (2018). See Supplementary Table S4 for details on the number of SNPs, number of monomorphic sites and models tested for each pair comparison.

We performed 50 independent *fastsimcoal2* runs per model per population pair. Each run consisted of 100,000 coalescent simulations and 40 expectation-maximization cycles for parameter optimization. We used a mutation rate of 1.0 x 10^−8^ based on Asteraceae EST sequence comparisons and fossil calibrations (Strasburg and Rieseberg 2008). We ranked the models based on the Kullback–Leibler information value which was estimated from the Akaike information criterion (AIC) scores of the best run per model. Here, the normalization of the difference between the AIC scores of a particular model and the best model in the set provides a measure of the degree of support for a particular model, namely model likelihood (*w_i_*) (Thomé and Carstens 2016). Since the use of linked-SNPs might lead to pseudo-replication issues when comparing models based on *fastsimcoal2* likelihood values (Bagley et al. 2017) and the SFS discards linkage information, we verified SNPs were largely unlinked by calculating linkage-disequilibrium in PLINK (data not shown).

As *fastsimcoal2* uses simulations to approximate the likelihood values, there is variance in the likelihood estimates. To test whether the best model significantly differs from alternative models with very low gene flow (2*Nm* = 0.01) but the same values at other parameters, we compared their likelihood distributions based on 100 expected SFS from 100,000 coalescent simulations per model (Bagley et al. 2017). If likelihood distributions overlap, there is no significant differences between the fit of both models (Meier et al. 2017). To obtain confidence intervals for all demographic parameters, we performed parametric bootstrapping. Given the parameter values of the best run of the best model, we simulated 100 SFS and re-estimated the parameter values from them. Each run consisted of 100,000 coalescent simulations and 30 expectation-maximization cycles. The parameter values of the best run of the best model were specified as the initial values of each bootstrapping run. We computed the 95% confidence intervals of all parameters with the *groupwiseMean* function of *rcompanion* R package (Mangiafico 2015).

### Is gene flow high enough to obscure a single origin scenario?

To better understand under what conditions gene flow can erode a signal of phylogenetic monophyly of each ecotype, we created a coalescent model to represent a single origin scenario of ecotypes within *Mathematica* v12.2 (Wolfram Research, Inc. 2020) (see Supplementary Methods S2 for full details). We assumed a species tree consisting of four populations, with two sets of sister taxa to represent populations of the same ecotype (Fig. 5A). The ancestor of the ecotypes splits at time *t2* in the past, and we can think of this split as an initial single origin of the Dune and Headland ecotypes. To represent two parapatric population pairs, each of these two ecotypes further split simultaneously at time, *t1*, in the past. We considered an instantaneous burst of migration from the populations of the ancestral ecotype into the populations of the derived ecotype at each locality by assuming that a fraction, *m*, of alleles in each derived ecotype population is replaced by migrant alleles from a parapatric population at time, *tm*, in the past. Considering a single lineage from each of the four populations starting at the present, we condition on the migrant status of the sampled lineages to calculate coalescent probabilities of gene tree topologies that result in a grouping in which the two populations of the derived ecotype are not monophyletic in the gene genealogy. These methods recapitulate recent more formal treatments of the probability of hemiplasy (non-monophyly despite a single mutational origin for a trait) under incomplete lineage sorting and introgression (Guerrero and Hahn 2018; Hibbins et al. 2020), though we have considered a scenario involving four populations to reflect the nature of parapatric pairs. Moreover, our emphasis is placed on the implications of gene flow for the original inference of the species tree itself, rather than how it pertains to the history of a selected locus of interest under an inferred phylogeny. Furthermore, we ran forward simulations of neutral polymorphism in *SLiM2* (Haller and Messer 2017) to further gain insight into the conditions in which gene flow can distort the phylogenetic topology. Instead of having an instantaneous unidirectional burst of migration, we considered a scenario of continuous and symmetrical migration. See Supplementary Methods S3-S5 for full description of the approach used.

Finally, although we cannot directly map where our observed data fall in the simulated parameter space, we can gain further confidence on the likelihood of phylogenetic distortion by considering divergence time estimates from *fastsimcoal2* in combination with the phylogenetic topology estimated in *IQ-TREE*. More specifically, we asked whether divergence times between populations from the same ecotype are deeper than between populations from different ecotypes. The estimation of divergence times in *fastsimcoal2* considers gene flow, thus if they are in accordance with relative node order of the *IQ-TREE* phylogeny (which is estimated without accounting for gene flow), it suggests that phylogenetic distortion within the system is unlikely. We thus compared the *fastsimcoal2* divergence times to the relative node order of the *IQ-TREE* phylogeny for four population pairs (D04-H05 and D05-H06; D14-H15 and D32-H12). We selected these pairs because they represent neighboring sister-taxa within the phylogeny.

## Acknowledgements

We are grateful to M.J. Wilkinson, H.N. Nguyen Vu and H.L. North for assisting with sample collection. We extend our thanks to R.K. Bagley for assisting with *fastsimcoal2*. We thank Ortiz-Barrientos laboratory members for insightful comments on previous versions of this manuscript. S. Chenoweth and M. Blows provided very useful feedback on M.E. James’ PhD dissertation. This work was supported by an Australian Research Council grant (DP190103039) to D.O. and J.E., and a University of Queensland Graduate School International Travel Award to M.E.J.

## Author contributions

M.E.J and D.O. conceived the project. M.E.J. and J.E. undertook sample collection. M.E.J. extracted DNA, prepared libraries, performed bioinformatics, and undertook the *IQ-TREE*, *fastSTRUCTURE*, *STRUCTURE* and *TreeMix* analyses. S.LA. assembled the reference genome. H.A. conducted the *fastsimcoal2* analyses. J.S.G. performed the coalescent modelling and *SLiM2* simulations with input from J.E. The paper was written by M.E.J. and D.O. with input from all authors. D.O. is the mentor and supervisor for the research program.

## Data and code availability

The data and code underlying this article are available in GitHub, at https://github.com/OB-lab/James_et_al._2021-MBE.

