## Supplementary for "Highly replicated evolution of parapatric ecotypes"

### Supplementary material Tables

#### Table S1. Pairwise $F_{ST}$ values for 21 *Senecio lautus* populations

| | D00 | D01 | D02 | D03 | D04 | D05 | D12 | D14 | D32 | H00 | H01 | H02 | H03 | H04 | H05 | H06 | H07 | H12 | H12A | H14 | H15 |
|---|---|---|---|---|---|---|---|---|---|---|---|---|---|---|---|---|---|---|---|---|---|
| **D00** | - | | | | | | | | | | | | | | | | | | | | |
| **D01** | 0.25 | - | | | | | | | | | | | | | | | | | | | |
| **D02** | 0.25 | 0.22 | - | | | | | | | | | | | | | | | | | | |
| **D03** | 0.27 | 0.22 | 0.20 | - | | | | | | | | | | | | | | | | | |
| **D04** | 0.29 | 0.25 | 0.26 | 0.28 | - | | | | | | | | | | | | | | | | |
| **D05** | 0.29 | 0.25 | 0.27 | 0.27 | 0.25 | - | | | | | | | | | | | | | | | |
| **D12** | 0.34 | 0.29 | 0.31 | 0.33 | 0.31 | 0.28 | - | | | | | | | | | | | | | | |
| **D14** | 0.34 | 0.28 | 0.29 | 0.31 | 0.29 | 0.26 | 0.32 | - | | | | | | | | | | | | | |
| **D32** | 0.34 | 0.30 | 0.33 | 0.34 | 0.32 | 0.30 | 0.30 | 0.32 | - | | | | | | | | | | | | |
| **H00** | 0.26 | 0.23 | 0.24 | 0.25 | 0.26 | 0.26 | 0.29 | 0.28 | 0.31 | - | | | | | | | | | | | |
| **H01** | 0.26 | 0.22 | 0.25 | 0.25 | 0.25 | 0.24 | 0.26 | 0.25 | 0.28 | 0.23 | - | | | | | | | | | | |
| **H02** | 0.26 | 0.21 | 0.21 | 0.20 | 0.27 | 0.27 | 0.31 | 0.30 | 0.33 | 0.25 | 0.25 | - | | | | | | | | | |
| **H03** | 0.33 | 0.29 | 0.31 | 0.32 | 0.30 | 0.27 | 0.28 | 0.30 | 0.30 | 0.29 | 0.25 | 0.31 | - | | | | | | | | |
| **H04** | 0.28 | 0.23 | 0.25 | 0.26 | 0.27 | 0.26 | 0.30 | 0.29 | 0.31 | 0.24 | 0.22 | 0.26 | 0.29 | - | | | | | | | |
| **H05** | 0.29 | 0.25 | 0.27 | 0.28 | 0.21 | 0.26 | 0.31 | 0.30 | 0.32 | 0.27 | 0.25 | 0.27 | 0.30 | 0.27 | - | | | | | | |
| **H06** | 0.30 | 0.27 | 0.28 | 0.29 | 0.27 | 0.21 | 0.30 | 0.28 | 0.31 | 0.27 | 0.24 | 0.28 | 0.28 | 0.27 | 0.28 | - | | | | | |
| **H07** | 0.31 | 0.27 | 0.28 | 0.29 | 0.28 | 0.24 | 0.28 | 0.28 | 0.30 | 0.27 | 0.24 | 0.29 | 0.27 | 0.27 | 0.28 | 0.26 | - | | | | |
| **H12** | 0.35 | 0.31 | 0.33 | 0.34 | 0.33 | 0.30 | 0.31 | 0.32 | 0.22 | 0.31 | 0.28 | 0.33 | 0.30 | 0.31 | 0.33 | 0.32 | 0.30 | - | | | |
| **H12A** | 0.34 | 0.30 | 0.33 | 0.33 | 0.32 | 0.30 | 0.29 | 0.31 | 0.20 | 0.31 | 0.27 | 0.32 | 0.29 | 0.31 | 0.32 | 0.32 | 0.30 | 0.22 | - | | |
| **H14** | 0.34 | 0.30 | 0.31 | 0.33 | 0.31 | 0.28 | 0.28 | 0.30 | 0.30 | 0.30 | 0.27 | 0.32 | 0.28 | 0.29 | 0.32 | 0.30 | 0.28 | 0.31 | 0.30 | - | |
| **H15** | 0.34 | 0.28 | 0.29 | 0.31 | 0.30 | 0.26 | 0.32 | 0.15 | 0.32 | 0.28 | 0.25 | 0.30 | 0.30 | 0.29 | 0.31 | 0.28 | 0.28 | 0.32 | 0.30 | 0.30 | - |

#### Table S2. Estimation of gene flow and other demographic parameters in *fastsimcoal2*

| Comparison | Populations[a] | Asize[b] | Pop1size[c] | Pop2size[d] | DivTime[e] | SecTime[f] | IsoTime[g] | 2*Nm*P1->P2[h] | 2*Nm*P2->P1[i] |
|---|---|---|---|---|---|---|---|---|---|
| Dune-Headland | D00-H00 | 100497 | 47926 | 134364 | 71945 | 18690 | NA | 0.2176 | 0.2830 |
| | D03-H02 | 88035 | 34637 | 152616 | 44190 | 15031 | NA | 0.1590 | 0.4722 |
| | D01-H01 | 72385 | 90270 | 159101 | 71918 | 13268 | NA | 0.6241 | 0.3671 |
| | D04-H05 | 84266 | 86607 | 107660 | 37211 | 7220 | 307 | **1.9889** | **1.2829** |
| | D05-H06 | 97653 | 131873 | 70970 | 67927 | 16810 | NA | 0.4049 | 0.4325 |
| | D12-H14 | 56510 | 211701 | 103102 | 110018 | 11783 | NA | 0.2188 | 0.1787 |
| | D14-H15 | 98848 | 39145 | 134647 | 61058 | 24349 | 1495 | 0.4569 | 0.7205 |
| | D32-H12 | 56568 | 661726 | 212041 | 38706 | 11290 | NA | **5.5694** | **5.2694** |
| Dune-Dune | D00-D02 | 97055 | 63843 | 113624 | 52711 | 6436 | NA | 0.3242 | 0.2617 |
| | D01-D03 | 99168 | 142624 | 51613 | 58652 | 23772 | NA | 0.3280 | 0.2119 |
| | D01-D04 | 92638 | 121936 | 74257 | 66970 | 11319 | NA | 0.2901 | 0.2200 |
| | D02-D03 | 93440 | 116800 | 48492 | 54857 | 23163 | NA | 0.3514 | 0.2029 |
| | D04-D05 | 77118 | 86082 | 115635 | 85983 | 19431 | 133 | 0.3751 | 0.2327 |
| | D05-D12 | 35322 | 103044 | 223346 | 128159 | 21689 | NA | 0.1562 | 0.2041 |
| | D12-D14 | 22223 | 259172 | 38450 | 118024 | 35758 | NA | 0.0991 | 0.1046 |
| | D14-D32 | 47348 | 27002 | 641721 | 56330 | 12595 | NA | 0.3179 | 0.1074 |
| Headland-Headland | H00-H02 | 97070 | 107516 | 94259 | 81290 | 9622 | NA | 0.3920 | 0.4012 |
| | H01-H04 | 78784 | 171269 | 86431 | 81443 | 19633 | NA | 0.2561 | 0.3261 |
| | H01-H05 | 61893 | 173061 | 89468 | 95145 | 17016 | NA | 0.2546 | 0.3116 |
| | H02-H04 | 78009 | 107213 | 91698 | 87055 | 20222 | NA | 0.2913 | 0.1768 |
| | H03-H07 | 57850 | 147099 | 125603 | 109197 | 12874 | NA | 0.1904 | 0.1921 |
| | H03-H14 | 63207 | 157400 | 119559 | 108683 | 9099 | NA | 0.2068 | 0.2012 |
| | H05-H06 | 84257 | 109077 | 88382 | 81710 | 10907 | NA | 0.2559 | 0.1953 |
| | H06-H07 | 67117 | 89121 | 141737 | 88627 | 14422 | NA | 0.2532 | 0.2387 |
| | H12-H12A | 52196 | 286574 | 322443 | 43928 | 14082 | NA | **3.7584** | **4.0315** |
| | H12-H15 | 46457 | 362929 | 51902 | 81116 | 9800 | NA | 0.1921 | 0.2501 |
| | H14-H15 | 46091 | 168257 | 72243 | 101092 | 18596 | NA | 0.1396 | 0.1181 |
| Allopatric | D03-D32 | 37805 | 51684 | 548897 | 81183 | 7366 | NA | 0.3340 | 0.2160 |
| | D03-H12 | 38444 | 61262 | 325186 | 104416 | 11011 | NA | 0.1982 | 0.1399 |
| | H02-H12 | 38546 | 76529 | 311384 | 113221 | 10998 | NA | 0.1988 | 0.1662 |

[a] The two populations used for each comparison (population 1 is on the left, and population 2 on the right)

[b] Ancestral effective population size (haploid numbers)

**Table S2 cont.**

[c] Effective population size of population 1 (haploid numbers)

[d] Effective population size of population 2 (haploid numbers)

[e] Divergence time between population 1 and population 2 (number of generations)

[f] Time since gene flow upon secondary contact (number of generations)

[g] Time since gene flow cessation (number of generations)

[h] Migration rate (2*Nm*) from population 1 to population 2 (forward in time)

[i] Migration rate (2*Nm*) from population 2 to population 1 (forward in time)

Values in bold represent 2*Nm* > 1

#### Table S3. 95% confidence intervals for gene flow estimates from 100 bootstrap runs inferred in *fastsimcoal2*

| Comparison | Populations[a] | 2*Nm*P1->P2min[b] | 2*Nm*P1->P2max[c] | 2*Nm*P2->P1min[d] | 2*Nm*P2->P1max[e] |
|---|---|---|---|---|---|
| Dune-Headland | D00-H00 | 0.2181 | 0.2281 | 0.2769 | 0.2865 |
| | D03-H02 | 0.1511 | 0.1611 | 0.4576 | 0.4758 |
| | D01-H01 | 0.5991 | 0.6174 | 0.3554 | 0.3655 |
| | **D04-H05** | **1.8266** | **1.9165** | **1.1682** | **1.2272** |
| | D05-H06 | 0.3924 | 0.4054 | 0.4172 | 0.4321 |
| | D12-H14 | 0.2174 | 0.2236 | 0.1775 | 0.1840 |
| | D14-H15 | 0.4201 | 0.4513 | 0.6712 | 0.7135 |
| | **D32-H12** | **4.9046** | **5.0810** | **5.2088** | **5.3999** |
| Dune-Dune | D00-D02 | 0.3186 | 0.3344 | 0.2576 | 0.2686 |
| | D01-D03 | 0.3183 | 0.3305 | 0.2040 | 0.2137 |
| | D01-D04 | 0.2818 | 0.2911 | 0.2148 | 0.2226 |
| | D02-D03 | 0.3417 | 0.3563 | 0.2000 | 0.2096 |
| | D04-D05 | 0.3787 | 0.3926 | 0.2367 | 0.2450 |
| | D05-D12 | 0.1533 | 0.1584 | 0.2020 | 0.2077 |
| | D12-D14 | 0.0976 | 0.1011 | 0.1037 | 0.1068 |
| | D14-D32 | 0.3109 | 0.3274 | 0.1050 | 0.1106 |
| Headland-Headland | H00-H02 | 0.3828 | 0.3952 | 0.3901 | 0.4038 |
| | H01-H04 | 0.2504 | 0.2574 | 0.3183 | 0.3290 |
| | H01-H05 | 0.2485 | 0.2558 | 0.3033 | 0.3139 |
| | H02-H04 | 0.2892 | 0.2978 | 0.1741 | 0.1802 |
| | H03-H07 | 0.1882 | 0.1945 | 0.1917 | 0.1975 |
| | H03-H14 | 0.2029 | 0.2089 | 0.1976 | 0.2038 |
| | H05-H06 | 0.2526 | 0.2610 | 0.1926 | 0.1992 |
| | H06-H07 | 0.2460 | 0.2537 | 0.2310 | 0.2382 |
| | **H12-H12A** | **3.6292** | **3.7754** | **3.9901** | **4.1278** |
| | H12-H15 | 0.1878 | 0.1954 | 0.2484 | 0.2574 |
| | H14-H15 | 0.1357 | 0.1415 | 0.1163 | 0.1202 |
| Allopatric | D03-D32 | 0.3176 | 0.3314 | 0.2136 | 0.2207 |
| | D03-H12 | 0.1928 | 0.2005 | 0.1384 | 0.1429 |
| | H02-H12 | 0.1972 | 0.2043 | 0.1644 | 0.1690 |

[a] The two populations used for each comparison; population 1 (*P1*) is on the left, and population 2 (*P2*) on the right

[b] The lower 95% CI for migration rate (2*Nm*) from *P1* to *P2*

[c] The upper 95% CI for migration rate (2*Nm*) from *P1* to *P2*

[d] The lower 95% CI for migration rate from *P2* to *P1*

[e] The upper 95% CI for migration rate from *P2* to *P1*

Migration rates are forward in time. Populations in bold represent 2*Nm* > 1

**Table S4. Parameter estimates for all tested models in *fastsimcoal2***

(see excel file for Supplementary Table S4)

#### Table S5. Sampling locations of the 23 *Senecio lautus* Dune and Headland populations

| Clade | Population code | Location | Ecotype | Pair | Coordinates[a] | N[b] |
|---|---|---|---|---|---|---|
| Eastern | D00 | QLD: Stradbroke Island | Dune | D00-H00 | S27° 31.153' E153° 30.189' | 62 |
| Eastern | H00 | QLD: Stradbroke Island | Headland | D00-H00 | S27° 26.140' E153° 32.749' | 63 |
| Eastern | D02 | QLD: Southport | Dune | - | S27° 56.846' E153° 25.736' | 62 |
| Eastern | D03 | NSW: Cabarita | Dune | **D03-H02** | S28° 19.794' E153° 34.264' | 61 |
| Eastern | H02 | NSW: Cabarita | Headland | **D03-H02** | S28° 21.013' E153° 34.676' | 61 |
| Eastern | H04 | NSW: Byron Bay | Headland | - | S28° 38.060' E153° 38.268' | 62 |
| Eastern | D01 | NSW: Lennox Head | Dune | D01-H01 | S28° 46.858' E153° 35.655' | 60 |
| Eastern | H01 | NSW: Lennox Head | Headland | D01-H01 | S28° 48.813' E153° 36.313' | 58 |
| Eastern | D04 | NSW: Coffs Harbour | Dune | **D04-H05** | S30° 18.946' E153° 08.142' | 62 |
| Eastern | H05 | NSW: Coffs Harbour | Headland | **D04-H05** | S30° 18.741' E153° 08.676' | 62 |
| Eastern | D05 | NSW: South West Rocks | Dune | **D05-H06** | S30° 53.027' E153° 04.037' | 62 |
| Eastern | H06 | NSW: South West Rocks | Headland | **D05-H06** | S30° 52.710' E153° 04.549' | 62 |
| South-eastern | H07 | NSW: Port Macquarie | Headland | - | S31° 28.526' E152° 56.219 | 60 |
| South-eastern | H03 | NSW: Kiama | Headland | - | S34° 40.301' E150° 51.704' | 63 |
| South-eastern | D12 | NSW: Bermagui | Dune | D12-H14 | S36° 28.346' E150° 03.581' | 62 |
| South-eastern | H14 | NSW: Green Cape | Headland | D12-H14 | S37° 15.748' E150° 02.991' | 62 |
| South-eastern | D32 | VIC: Cape Bridgewater | Dune | **D32-H12** | S38° 19.631' E141° 23.772' | 62 |
| South-eastern | H12 | VIC: Cape Bridgewater | Headland | **D32-H12** | S38° 22.728' E141° 22.018' | 63 |
| South-eastern | H12A | VIC: Cape Bridgewater | Intermediate | - | S38° 20.282' E141° 23.896' | 62 |
| South-eastern | D14 | TAS: Port Arthur | Dune | **D14-H15** | S43° 10.550' E147° 51.267' | 12 |
| South-eastern | H15 | TAS: Port Arthur | Headland | **D14-H15** | S43° 11.240' E147° 50.672' | 11 |
| Western | D35 | WA: Isthmus Hill | Dune | - | S35° 05.885' E117° 59.182' | 62 |
| Western | D09 | WA: Leeuwin-Naturaliste National Park | Dune | - | S33° 46.239' E114° 59.541' | 63 |

[a] Coordinates represent the mid-point of each population

[b] The final number of individuals after removing individuals with low genetic coverage

Parapatric pairs in bold are sister-taxa within the phylogeny. H12A is a population found within an ecotone between the Dune (D32) and Headland (H12) at this locality.

#### Table S6. Sequencing and alignment summary for *Senecio lautus* populations

| Population code | Mean # clean reads (range) | Mean % mapped reads (range) | % mapped reads properly paired (range) |
|---|---|---|---|
| D00 | 2,138,896 (971,466 - 3,506,240) | 94 (62 - 98) | 92 (61 - 96) |
| H00 | 3,075,580 (1,528,536 - 6,198,407) | 81 (16 - 97) | 79 (16 - 95) |
| D02 | 2,714,361 (895,858 - 5,258,091) | 80 (18 - 96) | 76 (17 - 94) |
| D03 | 3,160,935 (2,015,566 - 8,748,545) | 84 (21 - 97) | 78 (20 - 95) |
| H02 | 2,772,081 (1,408,465 - 4,192,718) | 85 (34 - 96) | 83 (33 - 94) |
| H04 | 3,176,210 (1,695,120 - 5,950,574) | 90 (72 - 97) | 79 (60 - 95) |
| D01 | 3,061,253 (1,318,262 - 4,548,766) | 96 (83 - 98) | 90 (72 - 96) |
| H01 | 2,770,561 (1,105,881 - 6,164,034) | 93 (42 - 98) | 91 (36 - 96) |
| D04 | 2,922,712 (2,146,253 - 3,718,635) | 91 (62 - 98) | 83 (61 - 96) |
| H05 | 2,866,233 (1,754,603 - 4,696,562) | 92 (71 - 97) | 85 (67 - 95) |
| D05 | 2,854,456 (1,554,814 - 4,156,601) | 93 (48 - 97) | 87 (44 - 94) |
| H06 | 2,112,573 (1,253,010 - 3,538,428) | 84 (37 - 97) | 82 (36 - 95) |
| H07 | 3,116,096 (1,646,581 - 10,437,355) | 82 (27 - 98) | 73 (21 - 96) |
| H03 | 2,795,169 (1,593,958 - 5,514,042) | 77 (15 - 97) | 76 (14 - 95) |
| D12 | 2,700,235 (1,448,045 - 5,032,607) | 90 (45 - 98) | 83 (39 - 94) |
| H14 | 3,033,007 (1,661,205 - 8,349,758) | 71 (11 - 96) | 67 (11 - 95) |
| D32 | 2,854,449 (1,517,908 - 5,609,011) | 79 (19 - 97) | 76 (17 - 95) |
| H12 | 2,892,473 (1,220,369 - 4,774,451) | 83 (34 - 97) | 80 (33 - 94) |
| H12A | 2,614,734 (1,509,934 - 8,120,979) | 85 (27 - 98) | 82 (27 - 95) |
| D14 | 2,894,283 (1,704,586 - 4,893,613) | 94 (75 - 98) | 85 (58 - 95) |
| H15 | 3,229,783 (1,823,447 - 4,958,055) | 90 (33 - 97) | 84 (29 - 94) |
| D35 | 2,987,725 (1,754,767 - 6,004,276) | 90 (62 - 98) | 78 (44 - 95) |
| D09 | 3,008,471 (1,794,627 - 4,826,686) | 67 (21 - 96) | 63 (20 - 92) |

The 19 individuals removed due to high missing data are not included.

#### Supplementary material Figures

##### Fig S1. Summary of *TreeMix* runs

**(A)** Maximum likelihood tree with no migration. **(B)** Residuals for the no migration tree. **(C)** Log-likelihoods for each model for 1-25 migration events. **(D)** Proportion of variance explained by each model for 1-25 migration events.

##### Fig S2. *TreeMix* migration events 1-10

Maximum likelihood tree with 10 migration events. Colored arrows denote the intensity and direction of migration events.

##### Fig S3. Frequency distribution of *f3*-statistics

Frequency distribution of *f3*-statistics calculated in *TreeMix* across all populations.

##### Fig S4. *fastSTRUCTURE* K=22, K=28 and marginal likelihoods

**(A)** Bayesian assignment of individuals to genetic clusters within *fastSTRUCTURE* for K=22 and K = 28. Each of the 1,319 individuals is depicted as a bar, with colors representing ancestry proportions to each cluster. Populations are ordered according to their geographic distribution along the coast. **(B)** Marginal likelihood values for successive K-values within *fastSTRUCTURE*. Red dashed lines denote the K-value that best explained the structure in the data (K = 22), as well as the K-value that maximized the marginal likelihood of the data (K = 28). **(C)** Change in marginal likelihoods from *fastSTRUCTURE* for successive K-values. Red dashed line denotes K = 23, higher K-values produce a negligible change in likelihood values.

##### Fig S5. *STRUCTURE* best K-values for the Dune-Headland pairs

*STRUCTURE* best K-values for the eight Dune-Headland replicate pairs based on the maximum value for $\Delta K$ (the second order rate of change in the log probability of data between successive K-values).

**Fig S6. Log-likelihood values for the ten demographic models tested in *fastsimcoal2* per pair**

**BM**: bidirectional migration. **M21**: migration from population 2 to 1. **M12**: migration from population 1 to 2. **BSC**: bidirectional migration after secondary contact. **SC21**: migration from population 2 to 1 after secondary contact. **SC12**: migration from population 1 to 2 after secondary contact. **EBM**: bidirectional migration after population splitting with cessation of gene flow. **ASC**: ancient bidirectional secondary contact. **PRSC**: population size reduction after bidirectional secondary contact. Note that the '**NM**: no migration model' is not depicted as its likelihood values are consistently very low in all comparisons.

**Fig S7. Likelihood values for testing whether gene flow is negligible**

**Max L**: maximum likelihood for the best run from the best model. **A**: fixed negligible gene flow (2*Nm* = 0.01) from population 2 (on the right) to population 1 (on the left). **B**: fixed negligible gene flow from population 1 to 2 (2*Nm* = 0.01). **C**: fixed negligible gene flow in both directions (2*Nm* = 0.01). The asterisk denotes the pair where at least one of the migration rates is not significantly different from 2*Nm* = 0.01.

**Fig S8. Forward population genetic simulations in *SLiM2***

**(A)** Schematic diagram representing the single origin scenario simulated in *SLiM2*. *T1* represents the time from the present to the second split, *T2* represents the time from the second split to the split of the ancestral population. Gene flow (denoted by dotted lines) occurs between populations within each locality. Light green and dark green circles represent populations from different ecotypes. **(B)** Logarithm of the ratio of mean Genealogical Sorting Index (*gsi*) calculated from simulations output in *SLiM2* under different combinations of branching times, population sizes and migration rates. Positive *gsi* ratio values (red) denote a stronger distorted signal of the single origin scenario, whereas negative *gsi* ratio values (blue) denote a stronger signal of the true single origin scenario. Overall, the patterns suggest that as the speed of diversification increases (from long to short internal branches), even small amounts of gene flow are likely to erode the true signal of a single origin where populations cluster by ecology and not by geography. This is because short internal branches are already likely to distort the phylogenetic signal due to high levels of ancestral polymorphism in each parapatric pair.

**Fig S1.**

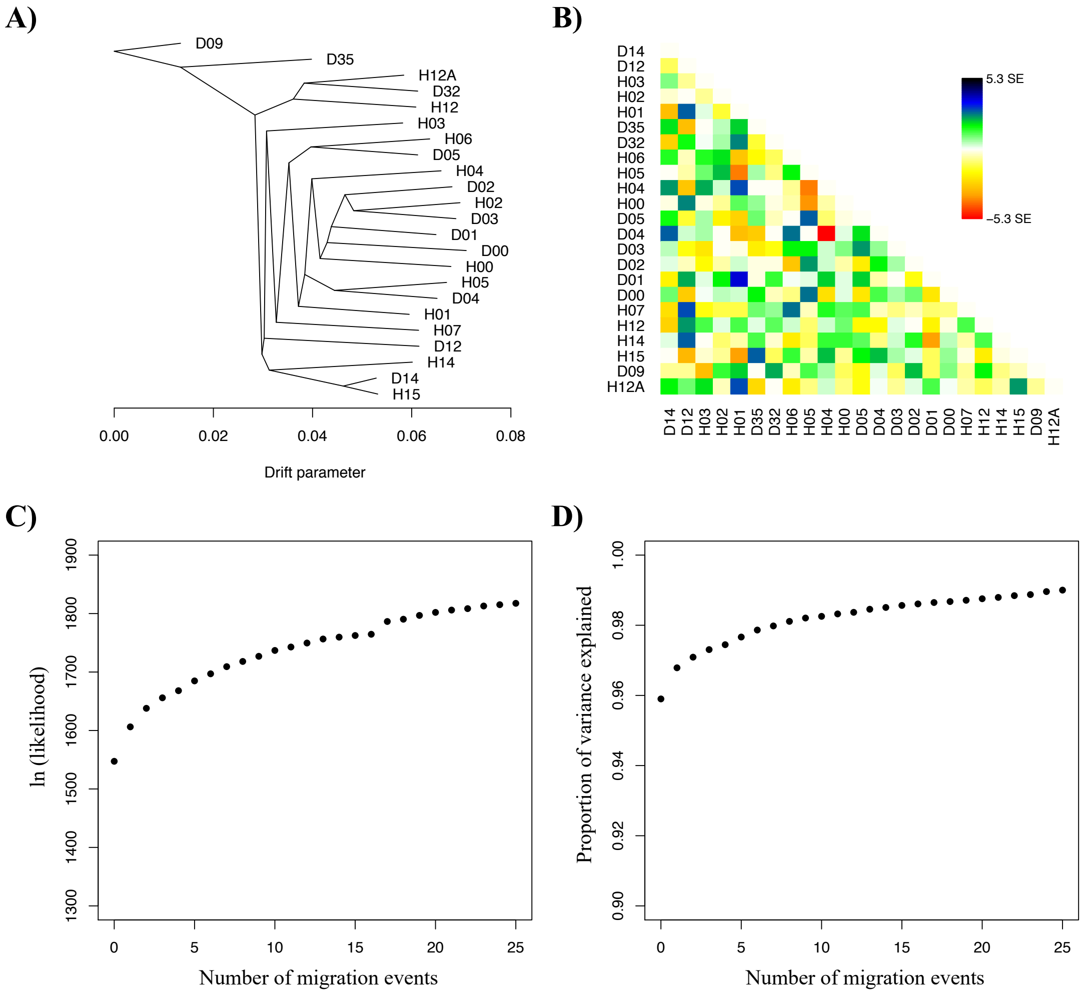
A)
B)
C)
D)
D09
D35
H12A
D32
H12
H03
H06
D05
H04
D02
H02
D03
D01
D00
H00
H05
D04
H01
H07
D12
H14
D14
H15
0.00
0.02
0.04
0.06
0.08
Drift parameter
5.3 SE
−5.3 SE
ln (likelihood)
Number of migration events
Proportion of variance explained
Number of migration events

**Fig S2.**

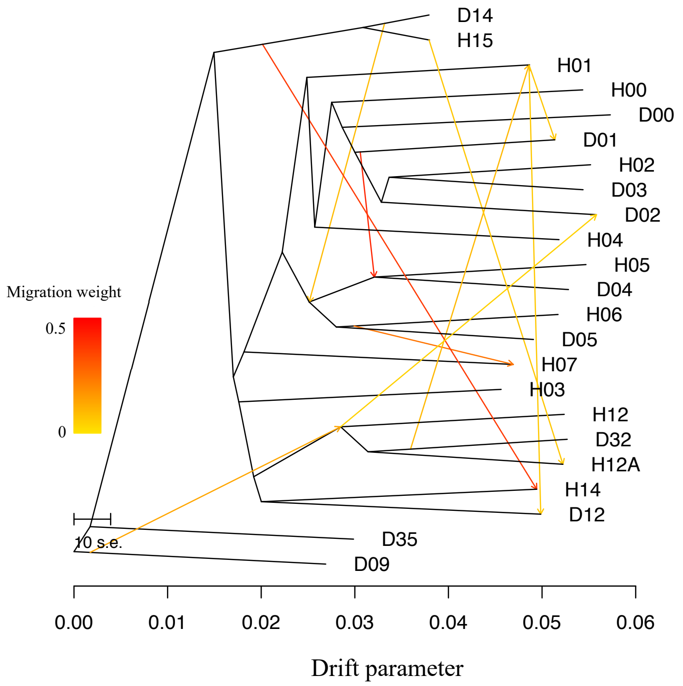
D14
H15
H01
H00
D00
D01
H02
D03
D02
H04
H05
D04
H06
D05
H07
H03
H12
D32
H12A
H14
D12
D35
D09
Migration weight
0.5
0
10 s.e.
0.00
0.01
0.02
0.03
0.04
0.05
0.06
Drift parameter

**Fig S3.**

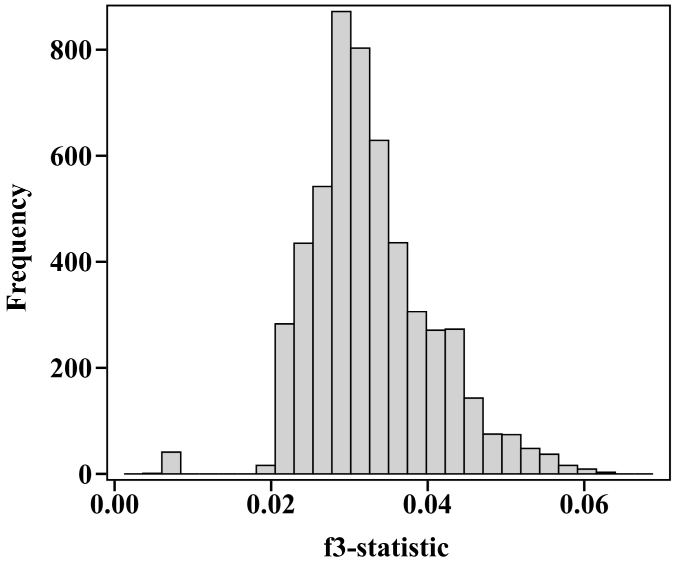

**Fig S4.**

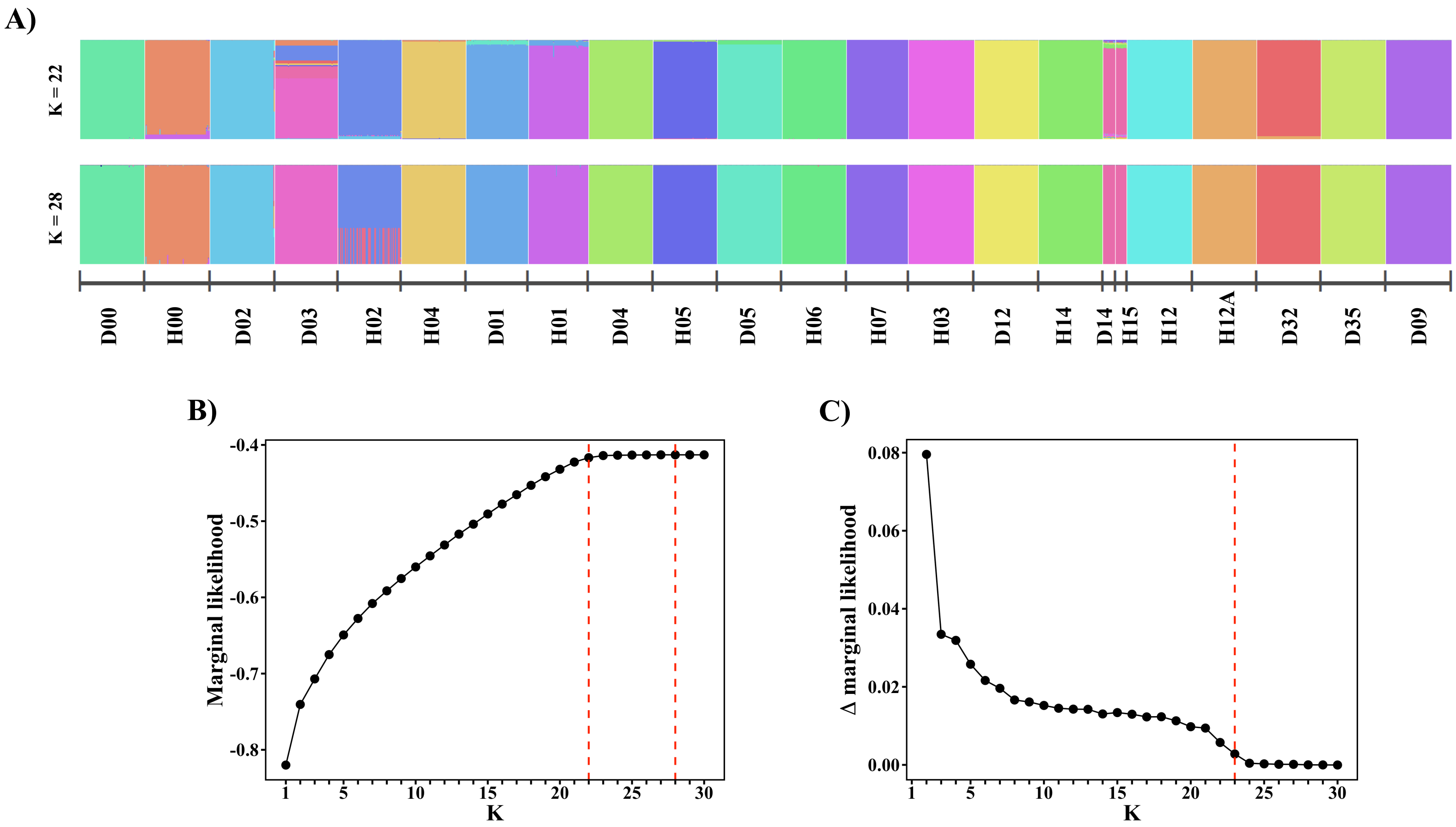

A)
K = 22
K = 28
D00
H00
D02
D03
H02
H04
D01
H01
D04
H05
D05
H06
H07
H03
D12
H14
D14
H15
H12
H12A
D32
D35
D09
B)
Marginal likelihood
K
C)
Δ marginal likelihood
K

**Fig S5.**

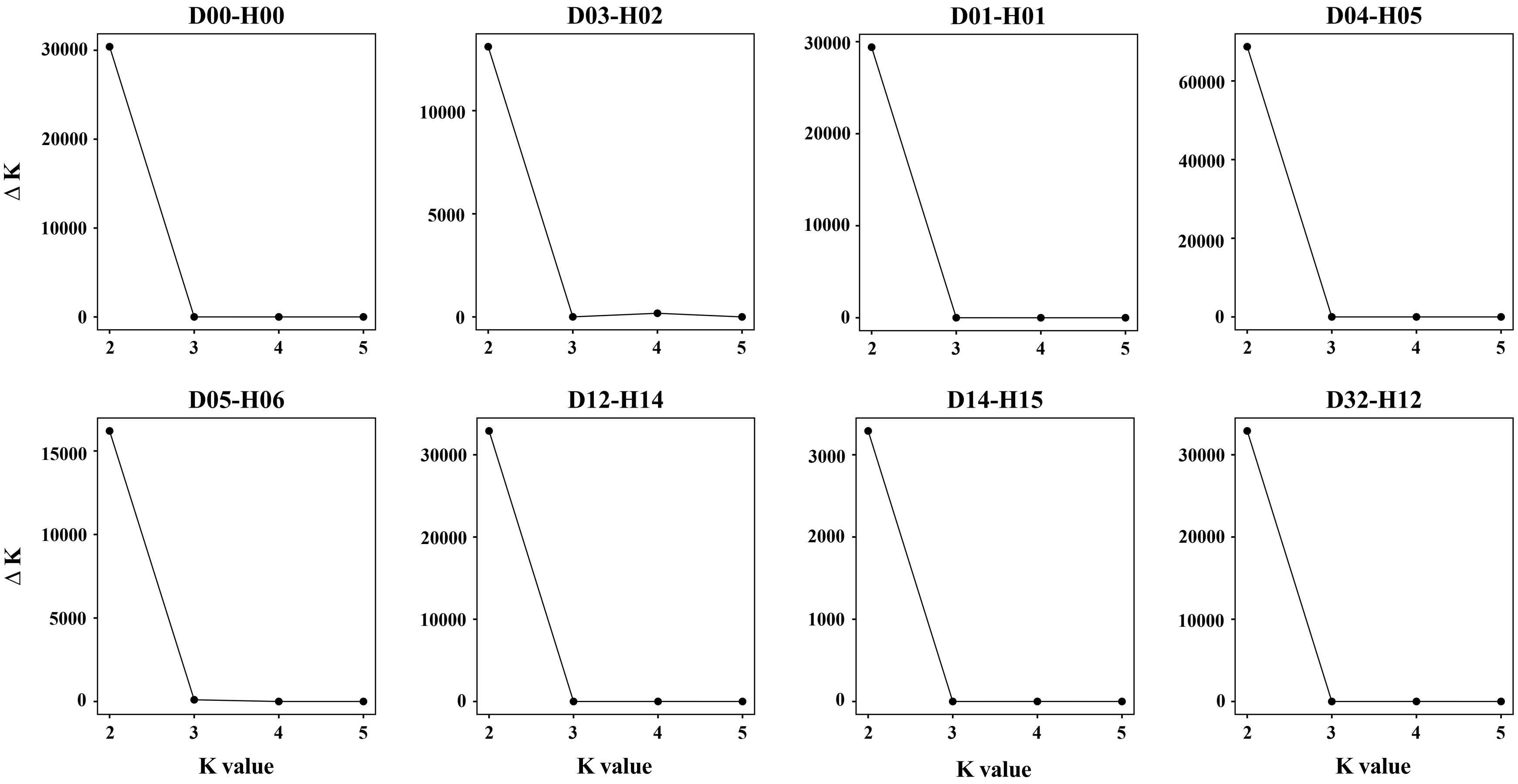
D00-H00
D03-H02
D01-H01
D04-H05
D05-H06
D12-H14
D14-H15
D32-H12
Δ K
K value

**Fig S6.**

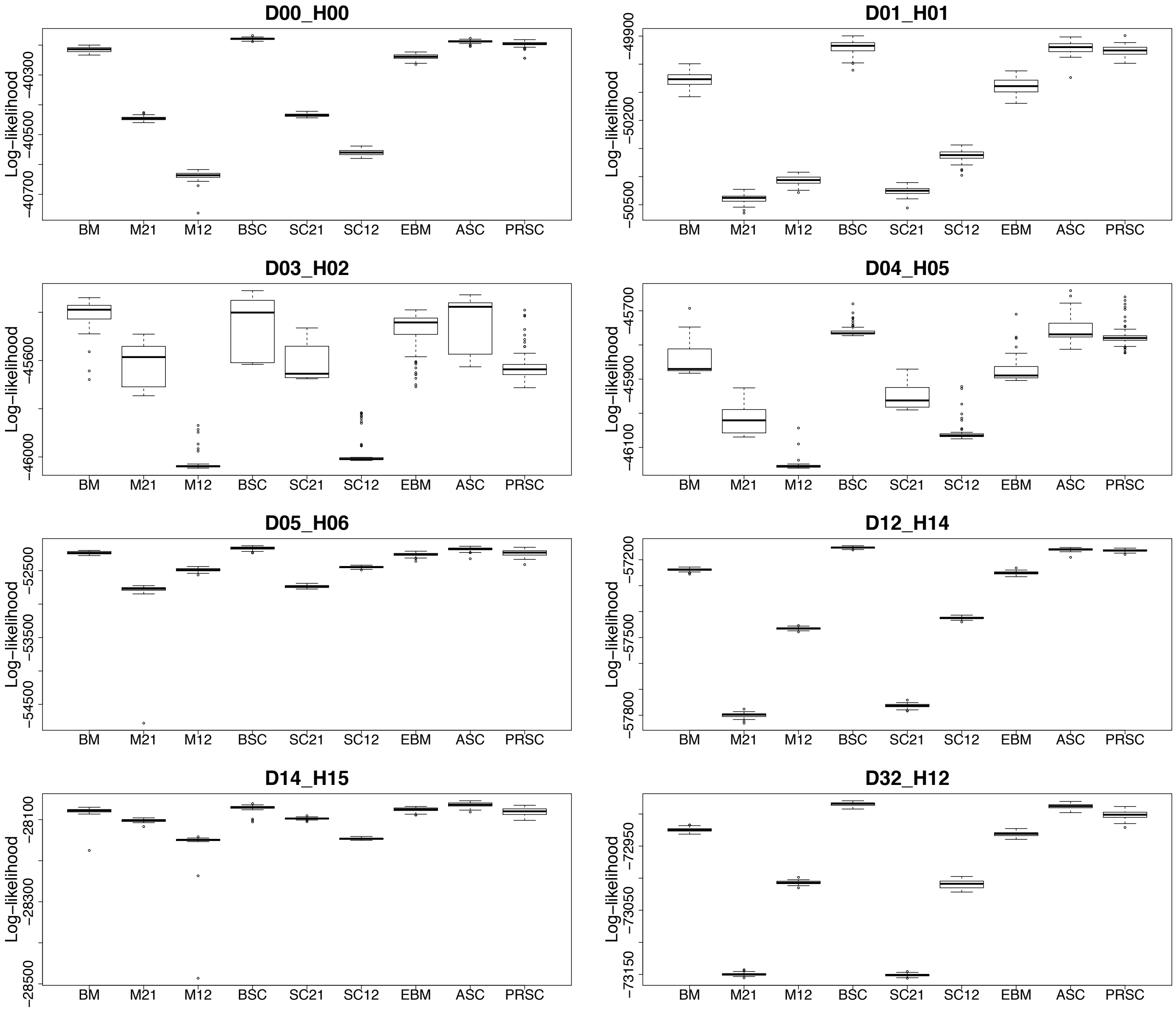
D00_H00
D01_H01
D03_H02
D04_H05
D05_H06
D12_H14
D14_H15
D32_H12
Log−likelihood
BM
M21
M12
BSC
SC21
SC12
EBM
ASC
PRSC

**Fig S6 cont.**

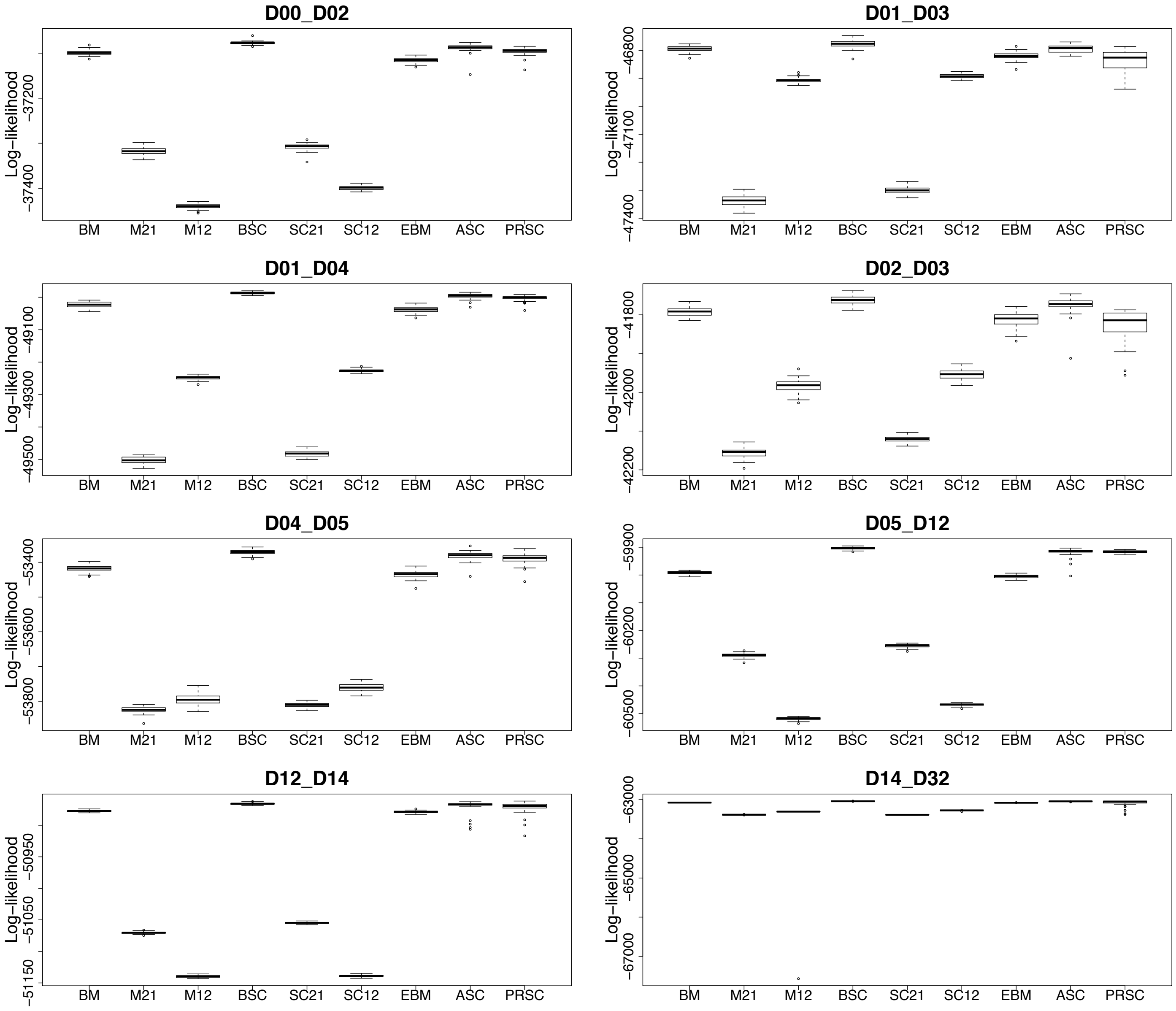

D00_D02
Log−likelihood
−37200
−37400
D01_D03
−46800
−47100
−47400
D01_D04
−49100
−49300
−49500
D02_D03
−41800
−42000
−42200
D04_D05
−53400
−53600
−53800
D05_D12
−59900
−60200
−60500
D12_D14
−50950
−51050
−51150
D14_D32
−63000
−65000
−67000
BM
M21
M12
BSC
SC21
SC12
EBM
ASC
PRSC

**Fig S6 cont.**

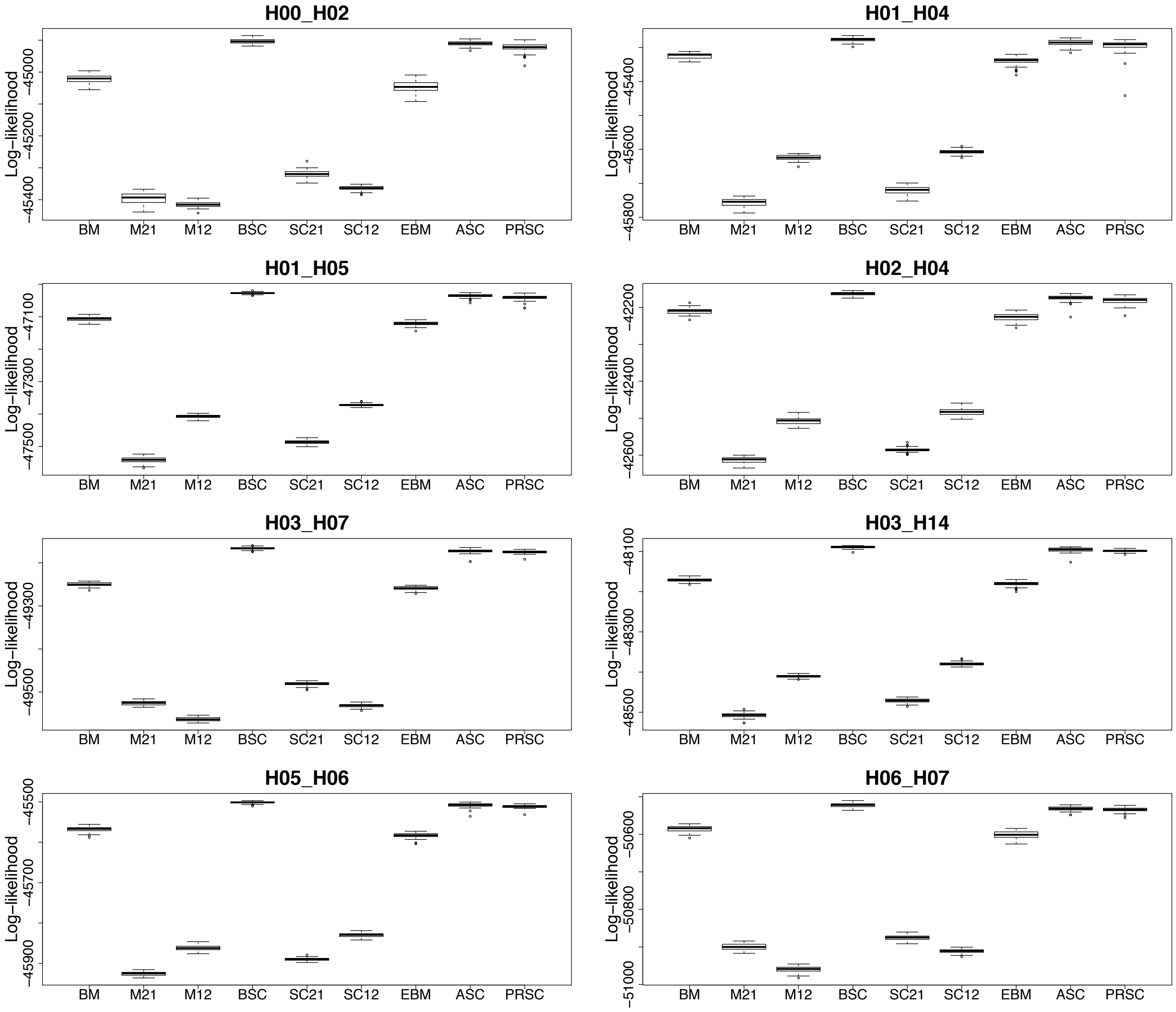
H00_H02
Log−likelihood
−45000
−45200
−45400
BM
M21
M12
BSC
SC21
SC12
EBM
ASC
PRSC
H01_H04
Log−likelihood
−45400
−45600
−45800
H01_H05
Log−likelihood
−47100
−47300
−47500
H02_H04
Log−likelihood
−42200
−42400
−42600
H03_H07
Log−likelihood
−49300
−49500
H03_H14
Log−likelihood
−48100
−48300
−48500
H05_H06
Log−likelihood
−45500
−45700
−45900
H06_H07
Log−likelihood
−50600
−50800
−51000

**Fig S6 cont.**

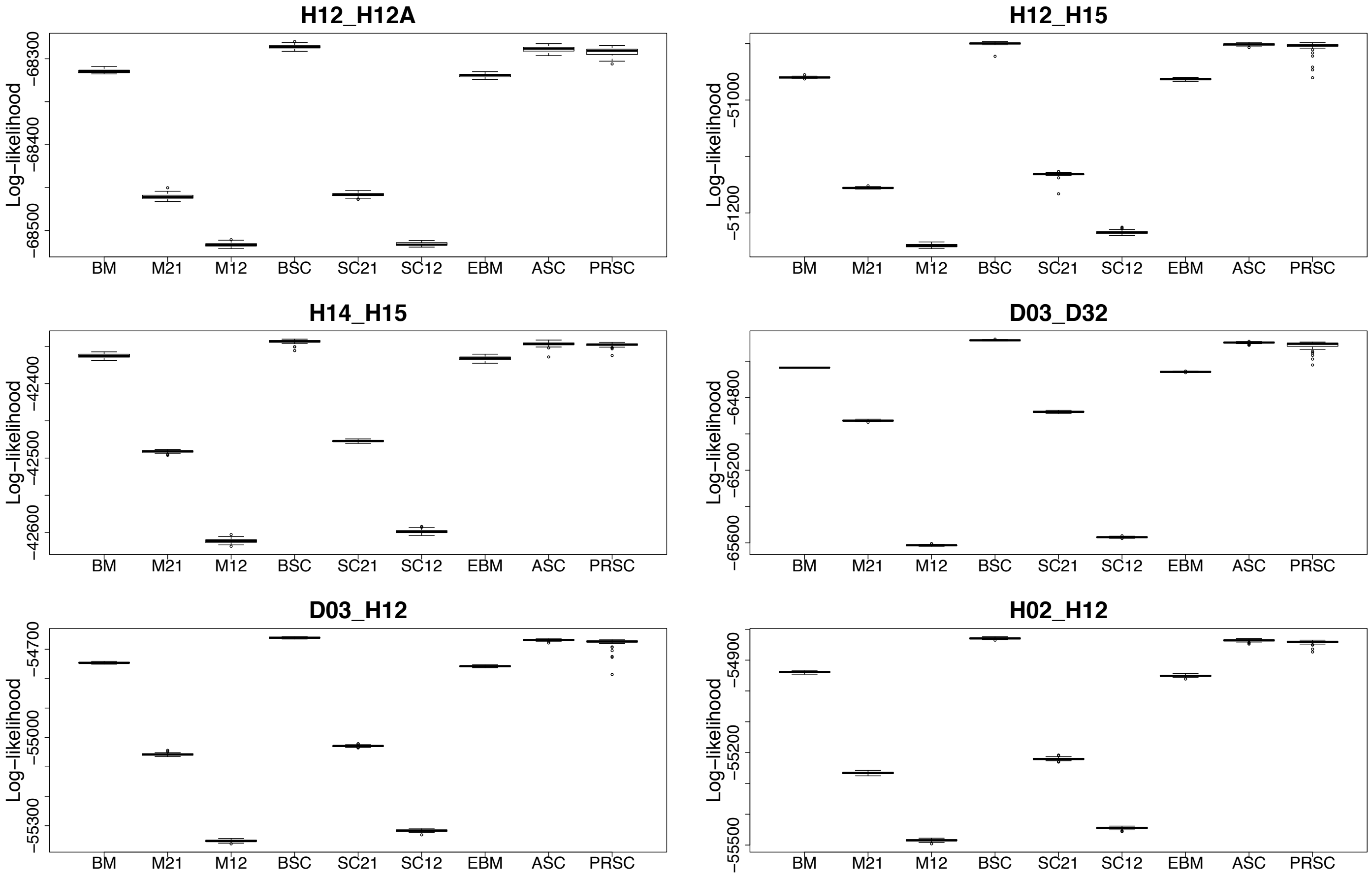
H12_H12A
H12_H15
H14_H15
D03_D32
D03_H12
H02_H12
Log−likelihood
BM
M21
M12
BSC
SC21
SC12
EBM
ASC
PRSC

**Fig S7.**

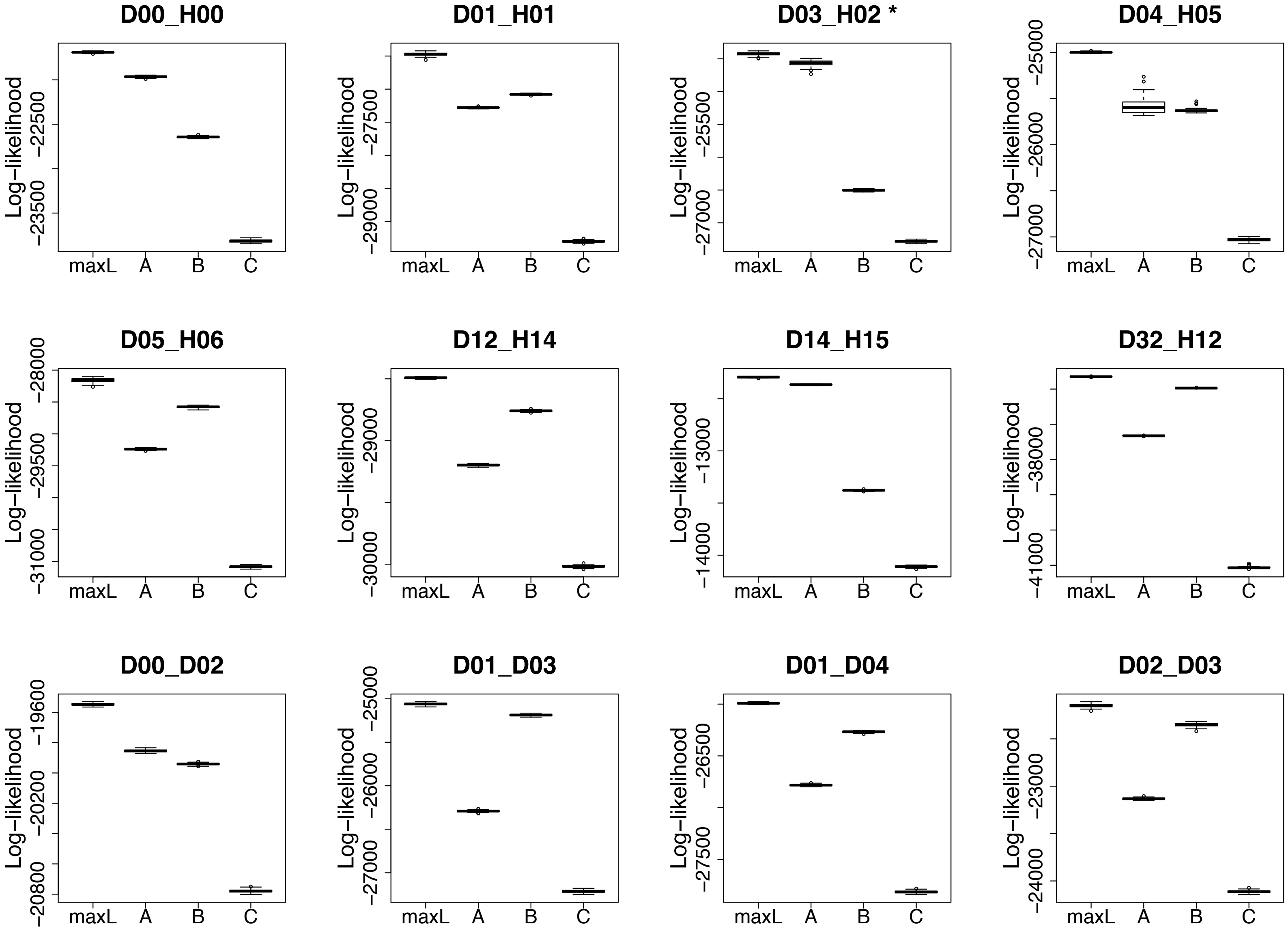
D00_H00
D01_H01
D03_H02 *
D04_H05
D05_H06
D12_H14
D14_H15
D32_H12
D00_D02
D01_D03
D01_D04
D02_D03
Log−likelihood
maxL
A
B
C

Fig S7 cont.

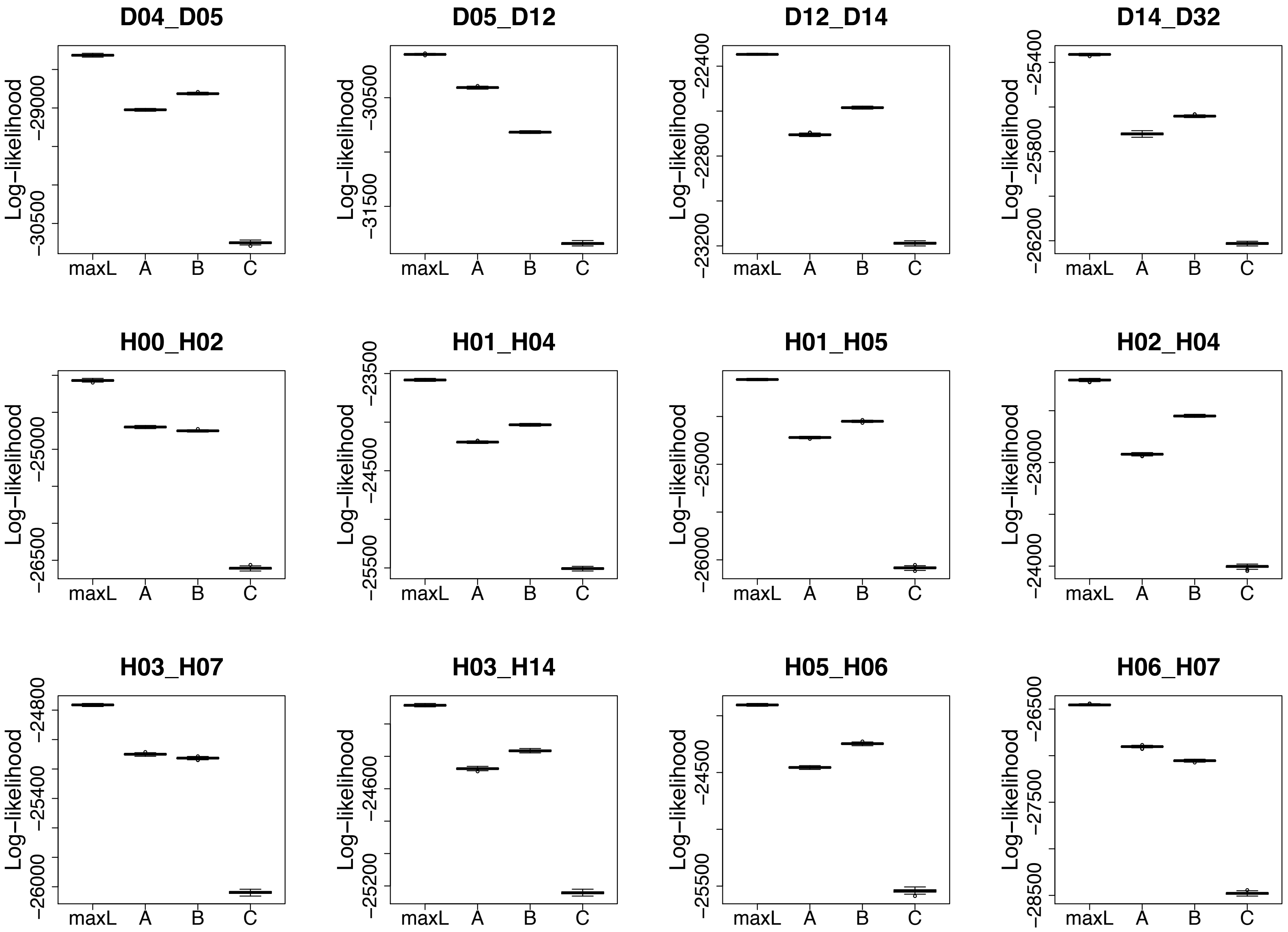

D04_D05
D05_D12
D12_D14
D14_D32
H00_H02
H01_H04
H01_H05
H02_H04
H03_H07
H03_H14
H05_H06
H06_H07
Log−likelihood
maxL
A
B
C

**Fig S7 cont.**

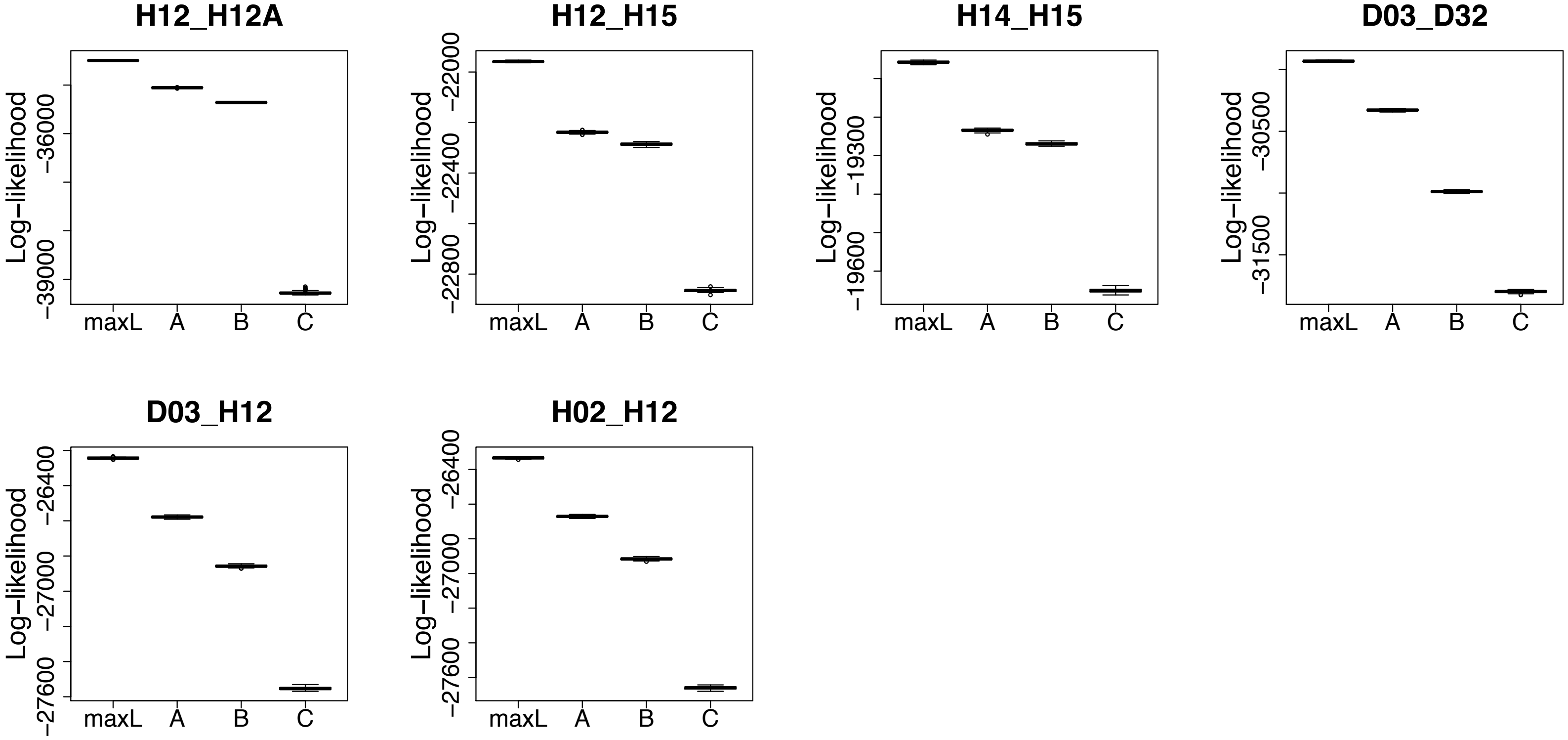
H12_H12A
Log−likelihood
−36000
−39000
maxL A B C
H12_H15
Log−likelihood
−22000
−22400
−22800
maxL A B C
H14_H15
Log−likelihood
−19300
−19600
maxL A B C
D03_D32
Log−likelihood
−30500
−31500
maxL A B C
D03_H12
Log−likelihood
−26400
−27000
−27600
maxL A B C
H02_H12
Log−likelihood
−26400
−27000
−27600
maxL A B C

**Fig S8.**

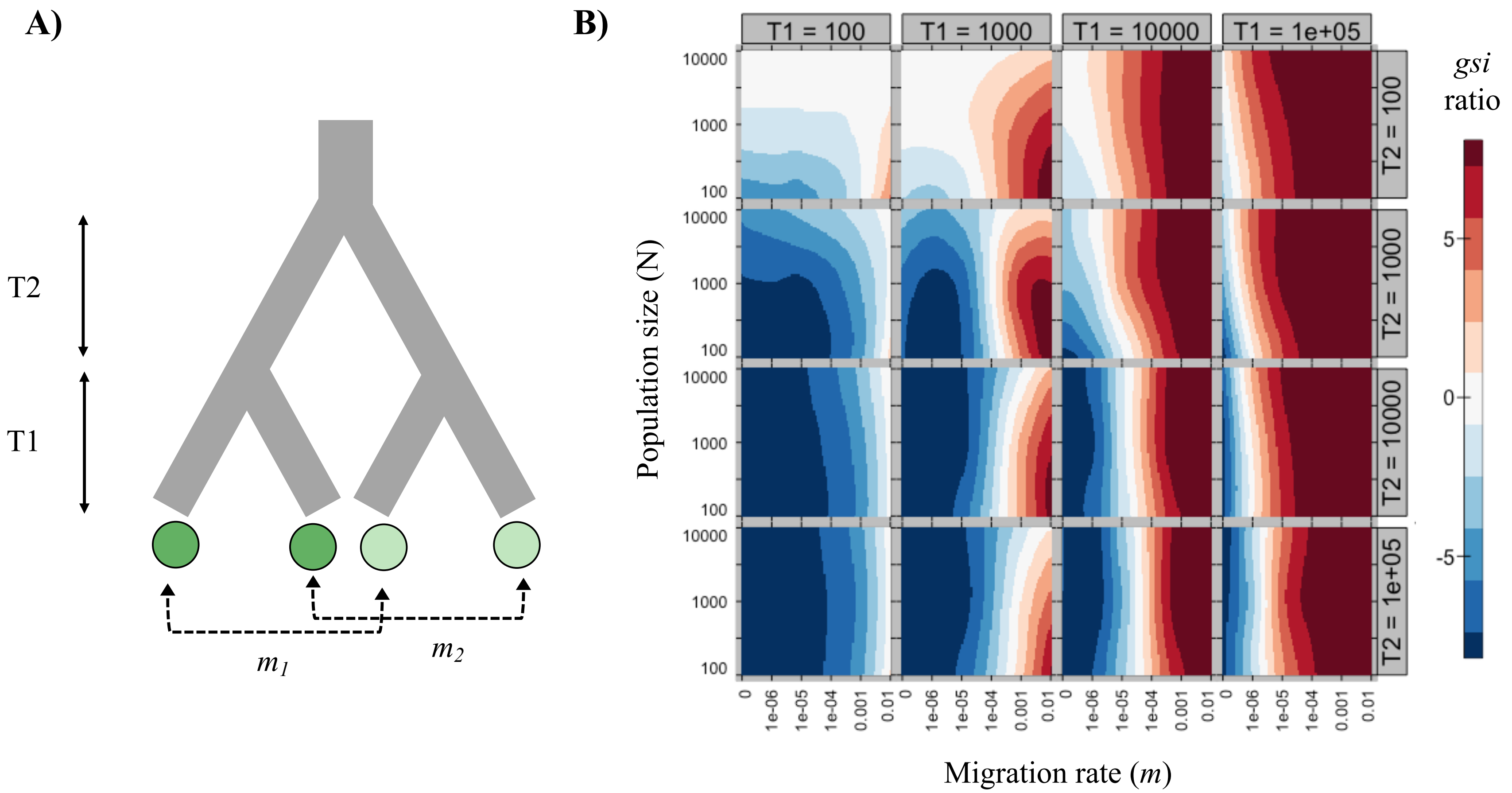
A)
T2
T1
$m_1$
$m_2$
B)
T1 = 100
T1 = 1000
T1 = 10000
T1 = 1e+05
T2 = 100
T2 = 1000
T2 = 10000
T2 = 1e+05
Population size (N)
10000
1000
100
Migration rate (m)
0
1e-06
1e-05
1e-04
0.001
0.01
gsi ratio
5
0
-5

#### Supplementary material Methods

##### S1 Methods. Estimation of monomorphic sites per pair

To estimate the number of monomorphic sites per pair we first calculated the number of RAD loci by using *PLINK* to thin for one SNP per RAD locus. The total read length of each RAD locus was (on average) 190bp (taking into account the length of the sequencing read after removal of barcodes/indexes). We used the following formula to calculate the number of monomorphic sites per pair:

*Monomorphic sites = (read length x number RAD loci) - number variable sites*

Here, we may be slightly overestimating the number of monomorphic sites as we are assuming all sites without a called SNP are monomorphic, although some could be actual variants that were not called due to not passing filtering requirements. Nevertheless, the parameter estimates (especially the migration rates) were robust to varying the number of monomorphic sites (data not shown).

### S2 Methods. Probabilities of gene flow distorting phylogeny topology

Consider a phylogeny of four populations, referred to as P1, P2, P3, P4. We have observed a species/population topology in which populations P3 and P4 are not sister. If we assume that the true species/population tree has the topology ((P1, P2), (P3, P4)), what is the probability that P3 and P4 are not sister under a model containing gene flow between P1 and P3 and between P2 and P4?

Alleles x1, x2, x3, x4 represent four lineages which are sampled from the respective populations at the present (t = 0). The lineages will be traced backwards in time, tracking coalescent events. The MRCA common ancestor of the sampled lineages is denoted x1,2,3,4. Other common ancestors are denoted likewise. Migration occurs from P1 to P3 and from P2 to P4 at tm. Migration occurs as an instantaneous burst and the parameter m represents the fraction of alleles in the recipient populations replaced by migrant alleles from the donor population. Divergence between P1 and P2 and between P3 and P4 occurs at time t1. The ancestor of P1 and P2 diverges from the ancestor of P3 and P4 at time t2.

The approach below will sum coalescent probabilities of all mutually exclusive ways in which (3,4) are not closest relatives. The probability of each scenario will be assigned to a variable and these will be summed at the end. Probabilities will be conditioned on whether x3 and x4 are descended from migrant alleles.

---

**Condition on x3 being descended from a migrant, x4 not descended from a migrant**

S1. x1 and x3 coalesce between tm and t1

In[1]:=
```
S1 = 1 - E^(-(t1 - tm))
```

Out[1]= $1 - e^{-t1+tm}$

S2. x1 and x3 don't coalesce between tm and t1. In the ancestral lineage containing exclusively x1, x2, and x3, a single coalescent event occurs between t1 and t2. That coalescent event is either (x3 with x1) or (x3 with x2).

In[2]:=
```
t = t2 - t1
OneCoal = Integrate[3 E^(-3 s) (E^( -(t - s))), {s, 0, (t)}]
S2 = (1 - S1) OneCoal (2/3)
```

Out[2]= $-t1 + t2$

Out[3]= $\frac{3}{2} e^{t1-3\,t2} \left(-e^{2\,t1} + e^{2\,t2}\right)$

Out[4]= $e^{-3\,t2+tm} \left(-e^{2\,t1} + e^{2\,t2}\right)$

S3. x1 and x3 don't coalesce between tm and t1. In the ancestral lineage containing exclusively x1, x2, and x3, a single coalescent event occurs between t1 and t2. That coalescent event is (x1 with x2). In the ancestral lineage containing x1,2, x3, and x4, the first coalescent event that occurs is either (x1,2 with x3) or (x1,2 with x4).

In[5]:=
```
S3 = (1 - S1) OneCoal (1/3) (2/3)
```

Out[5]= $\frac{1}{3} e^{-3\,t2+tm} \left(-e^{2\,t1} + e^{2\,t2}\right)$

S4. x1 and x3 don't coalesce between tm and t1. In the ancestral lineage containing exclusively x1, x2, and x3, two coalescent events occur, the first is (x1 with x2) and the second is (x1,2 with x3)

In[6]:=
```
TwoCoal = Integrate[3 E^(-3 s) (1 - E^( -(t - s))), {s, 0, t}]
S4 = (1 - S1) TwoCoal (1/3)
```

Out[6]= $\frac{1}{2}\left(2 + e^{3\,t1-3\,t2} - 3\,e^{t1-t2}\right)$

Out[7]= $\frac{1}{6}\,e^{-t1+tm}\left(2 + e^{3\,t1-3\,t2} - 3\,e^{t1-t2}\right)$

S5. x1 and x3 don't coalesce between tm and t1. In the ancestral lineage containing exclusively x1, x2, and x3, zero coalescent events occur between t1 and t2. In the ancestral lineage containing x1, x2, x3, and x4, the first coalescent event that occurs is one of (x3 with x1), (x3 with x2), (x1 with x4), (x2 with x4).

In[8]:=
```
S5 = (1 - S1) E^(-3 (t2 - t1)) (4/6)
```

Out[8]= $\frac{2}{3}\,e^{-t1-3\,(-t1+t2)+tm}$

S6. x1 and x3 don't coalesce between tm and t1. In the ancestral lineage containing exclusively x1, x2, and x3, zero coalescent events occur between t1 and t2. In the ancestral lineage containing x1, x2, x3, and x4, the first coalescent event that occurs is (x1 with x2). The next coalescent event to occur is either (x1,2 with x3) or (x1,2 with x4).

In[9]:=
```
S6 = (1 - S1) E^(-3 (t2 - t1)) (1/6) (2/3)
```

Out[9]= $\frac{1}{9}\,e^{-t1-3\,(-t1+t2)+tm}$

Putting this together, the probability that x3 is descended from a migrant, x4 is not descended from a migrant, and that conditional on this scenario x3 and x4 do not appear as sister in the phylogeny is:

In[10]:=
```
Prob1 = m (1 - m) (S1 + S2 + S3 + S4 + S5 + S6)
```

Out[10]= $\left(1 - e^{-t1+tm} + \frac{7}{9}\,e^{-t1-3\,(-t1+t2)+tm} + \frac{1}{6}\,e^{-t1+tm}\left(2 + e^{3\,t1-3\,t2} - 3\,e^{t1-t2}\right) + \frac{4}{3}\,e^{-3\,t2+tm}\left(-e^{2\,t1} + e^{2\,t2}\right)\right)(1-m)\,m$

---

**Condition on x4 being descended from a migrant, x3 not descended from a migrant**

By symmetry, the probability that x4 is descended from a migrant, x3 is not descended from a migrant, and that conditional on this scenario x3 and x4 not being each other's closest relatives is:

In[11]:=
```
Prob2 = Prob1
```

Out[11]= $\left(1 - e^{-t1+tm} + \frac{7}{9}\,e^{-t1-3\,(-t1+t2)+tm} + \frac{1}{6}\,e^{-t1+tm}\left(2 + e^{3\,t1-3\,t2} - 3\,e^{t1-t2}\right) + \frac{4}{3}\,e^{-3\,t2+tm}\left(-e^{2\,t1} + e^{2\,t2}\right)\right)(1-m)\,m$

---

**Condition on neither x3 nor x4 being descended from migrants**

S7. x3 and x4 don't coalesce between t1 and t2. x1 and x2 don't coalesce between t1 and t2. In the ancestral lineage containing x1, x2, x3, and x4, the first coalescent event is one of (x3 with x1), (x3 with x2), (x1 with x4), (x2 with x4).

In[12]:=
```
S7 = (E^(-(t2 - t1)))^2 (4/6)
```

Out[12]= $\frac{2}{3}\,e^{2\,t1-2\,t2}$

S8. x3 and x4 don't coalesce between t1 and t2. x1 and x2 don't coalesce between t1 and t2. In the ancestral lineage containing x1, x2, x3, and x4, the first coalescent event is (x1 with x2). The next coalescent event is one of (x1,2 with x3), (x1,2, x4).

In[13]:= S8 = (E^(−(t2 − t1)))^2 (1/6) (2/3)

Out[13]= $\frac{1}{9} e^{2\,t1-2\,t2}$

S9. x3 and x4 don't coalesce between t1 and t2. x1 and x2 coalesce between t1 and t2. In the ancestral lineage containing x1,2, x3, and x4, the first coalescent event is one of (x1,2 with x3), (x1,2, x4).

In[14]:= S9 = E^(−(t2 − t1)) (1 − E^(−(t2 − t1))) (2/3)

Out[14]= $\frac{2}{3} e^{t1-t2} \left(1 - e^{t1-t2}\right)$

Putting this together, the probability that neither x3 nor x4 are descended from migrants, and conditional on this x3 and x4 not being each other's closest relatives is:

In[15]:= Prob3 = (1 − m)^2 (S7 + S8 + S9)

Out[15]= $\left(\frac{7}{9} e^{2\,t1-2\,t2} + \frac{2}{3} e^{t1-t2} \left(1 - e^{t1-t2}\right)\right) (1-m)^2$

---

**Condition on both x3 and x4 being descended from migrants.**

S10. x1 and x3 coalesce between tm and t1. x2 and x4 do not coalesce between tm and t1.

In[16]:= S10 = (1 − E^(−(t1 − tm))) E^(−(t1 − tm))

Out[16]= $e^{-t1+tm} \left(1 - e^{-t1+tm}\right)$

S11. x1 and x3 do not coalesce between tm and t1. x2 and x4 coalesce between tm and t1. By symmetry, this is the same as above

In[17]:= S11 = S10

Out[17]= $e^{-t1+tm} \left(1 - e^{-t1+tm}\right)$

S12. x1 and x3 coalesce between tm and t1. x2 and x4 coalesce between tm and t1.

In[18]:= S12 = (1 − E^(−(t1 − tm)))^2

Out[18]= $\left(1 - e^{-t1+tm}\right)^2$

S13. x1 and x3 do not coalesce between tm and t1. x2 and x4 do not coalesce between tm and t1. In the ancestral lineage containing x1, x2, x3, and x4, the first coalescent event is one of (x3 with x1), (x3 with x2), (x1 with x4), (x2 with x4).

In[19]:= S13 = (E^(−(t1 − tm)))^2 (4/6)

Out[19]= $\frac{2}{3} e^{-2\,t1+2\,tm}$

S14. x1 and x3 do not coalesce between tm and t1. x2 and x4 do not coalesce between tm and t1. In the ancestral lineage containing x1, x2, x3, and x4, the first coalescent event is (x1 with x2). The next coalescent event is either (x1,2 with x3) or (x1,2 with x4).

In[20]:= S14 = (E^(−(t1 − tm)))^2 (1/6) (2/3)

Out[20]= $\frac{1}{9} e^{-2\,t1+2\,tm}$

Putting this together, the probability of x3 and x4 both being descended from migrant alleles, and conditional on this x3 and x4 not being each other's closest relatives is:

In[21]:= Prob4 = m^2 (S10 + S11 + S12 + S13 + S14)

Out[21]= $\left(\frac{7}{9} e^{-2\, t1+2\, tm} + 2\, e^{-t1+tm} \left(1 - e^{-t1+tm}\right) + \left(1 - e^{-t1+tm}\right)^2\right) m^2$

---

**Then from the law of total probability, we have that the probability of x3 and x4 not being each other's closest relatives is:**

In[22]:= P = Prob1 + Prob2 + Prob3 + Prob4

Out[22]= $\left(\frac{7}{9} e^{2\, t1-2\, t2} + \frac{2}{3} e^{t1-t2} \left(1 - e^{t1-t2}\right)\right) (1 - m)^2 +$
$2 \left(1 - e^{-t1+tm} + \frac{7}{9} e^{-t1-3\,(-t1+t2)+tm} + \frac{1}{6} e^{-t1+tm} \left(2 + e^{3\, t1-3\, t2} - 3\, e^{t1-t2}\right) + \frac{4}{3} e^{-3\, t2+tm} \left(-e^{2\, t1} + e^{2\, t2}\right)\right) (1 - m)\, m +$
$\left(\frac{7}{9} e^{-2\, t1+2\, tm} + 2\, e^{-t1+tm} \left(1 - e^{-t1+tm}\right) + \left(1 - e^{-t1+tm}\right)^2\right) m^2$

---

**Visuals**

In[23]:= PPlot1 = P /. t2 → (t3 + t1) /. t1 → 1 /. tm → .1

PPlot2 = P /. t2 → t1 + 1 /. t1 → tm + t4 /. tm → .1

Out[23]= $\left(\frac{7}{9} e^{2-2\,(1+t3)} + \frac{2}{3} e^{-t3} \left(1 - e^{-t3}\right)\right) (1 - m)^2 +$
$2 \left(0.59343 + \frac{7}{9} e^{-0.9-3\, t3} + \frac{4}{3} e^{0.1-3\,(1+t3)} \left(-e^{2} + e^{2\,(1+t3)}\right) + 0.0677616 \left(2 - 3\, e^{-t3} + e^{3-3\,(1+t3)}\right)\right) (1 - m)\, m +$
$0.963267\, m^2$

Out[24]= $\left(\frac{2\left(1 - \frac{1}{e}\right)}{3\, e} + \frac{7}{9} e^{2\,(0.1+t4)-2\,(1.1+t4)}\right) (1 - m)^2 +$
$2 \left(1 + \frac{7\, e^{-3-t4}}{9} - e^{-t4} + \frac{4}{3} e^{0.1-3\,(1.1+t4)} \left(-e^{2\,(0.1+t4)} + e^{2\,(1.1+t4)}\right) + \frac{1}{6} e^{-t4} \left(2 - \frac{3}{e} + e^{3\,(0.1+t4)-3\,(1.1+t4)}\right)\right) (1 - m)\, m +$
$\left(\frac{7}{9} e^{0.2-2\,(0.1+t4)} + 2\, e^{-t4} \left(1 - e^{-t4}\right) + \left(1 - e^{-t4}\right)^2\right) m^2$

In[25]:=

```
ContourPlot [PPlot1 , {m, 0, .1}, {t3, ((1/50)), 1}, PlotLegends → Automatic ,
  Axes → False , Frame → {True, True, False , False }, FrameLabel → {m, t2 - t1},
  LabelStyle → Directive [FontSize → 16], FrameTicksStyle → Directive [FontSize → 14]]
ContourPlot [PPlot2 , {m, 0, .1}, {t4, ((1/50)), 1}, PlotLegends → Automatic ,
  Axes → False , Frame → {True, True, False , False }, FrameLabel → {m , t1 - tm },
  FrameTicksStyle → Directive [FontSize → 14], LabelStyle → Directive [FontSize → 16]]
```

Out[25]=

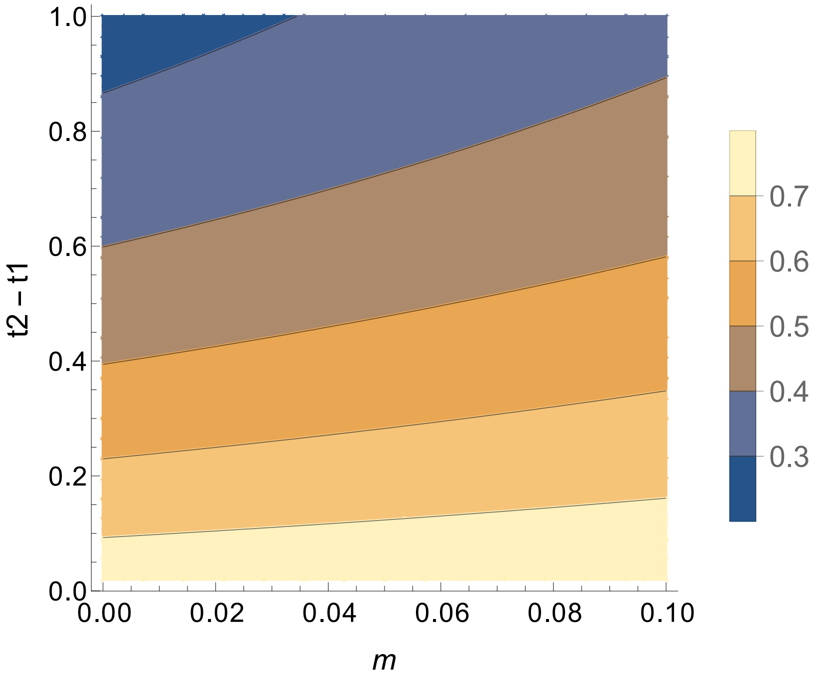


Out[26]=

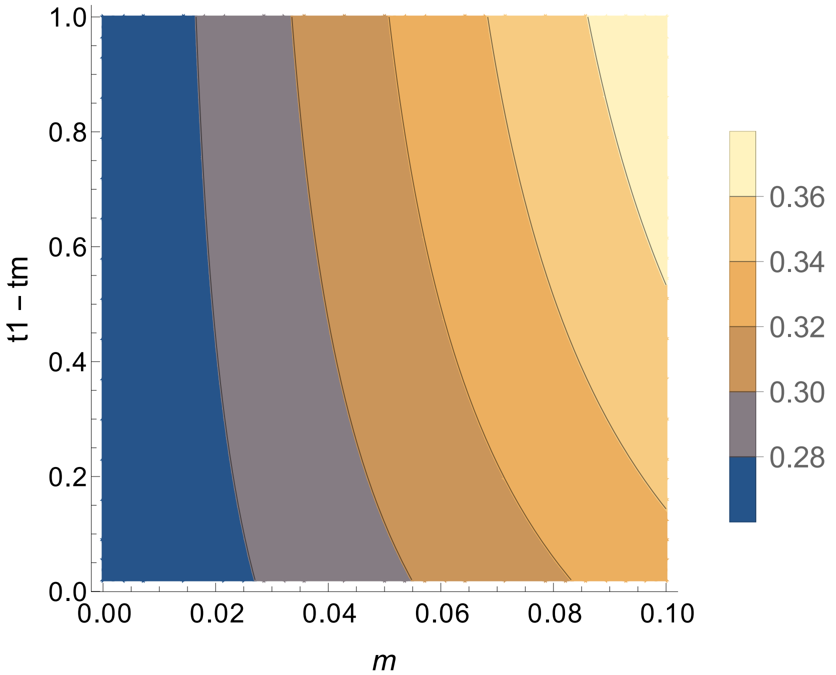

#### S3 Methods. Forward simulations in *SLiM2*

*SLiM2* (Haller and Messer 2017) simulates diploid genomes using a Wright-Fisher model, and tracks derived mutations within simulated genomes. As with the coalescent modelling, we mimicked a model of a single-origin scenario. Instead of having an instantaneous burst of migration, we considered a scenario of continuous symmetrical migration, where one population of one ecotype exchanges genes with the other ecotype within each locality (representing one parapatric pair at *location 1*, and this also occurs at *location 2* to represent the other parapatric pair, see Supplementary Fig. S8A). We varied the following parameters: population size, migration rate, time from the present (into the past) to the second split (*T1*), and time from the second split to the split of the ancestral population (*T2*). Note that the labels for *T1* and *T2* in the script are reversed (see Supplementary Methods S4). In addition, an outgroup population was retained after the first split to construct a rooted tree. Each model used a heuristic burn-in period of 10*N* generations to reach mutation-drift balance in the ancestral population. After each simulation, 30 individuals per population were sampled and output in a VCF file. We ran 30 replicate runs for each parameter combination. See Supplementary Methods S4 for *SLiM2* code.

We calculated a distance matrix between individual genotypes and constructed a rooted neighbor-joining tree within the *ape* R package (Paradis et al. 2004; Popescu et al. 2012). We calculated the genealogical sorting index (*gsi*; Cummings et al. 2008) to measure of how monophyletic an arbitrary set of tips are on a tree (see Supplementary Methods S5 for code). If all the tips of the phylogeny form a monophyletic group, the *gsi* will be 1, whereas if the tips are dispersed throughout the phylogeny, then the *gsi* will be closer to 0. We calculated the *gsi* for four sets of tips on each tree: for both populations from *location 1*, for both populations from *location* 2, for all 'Dunes', and for all 'Headlands'. We took the average of the first two (same location), and the average of the second two (same ecotype), then the natural log of the ratio of these two *gsi* values. When this *gsi* ratio is large and positive, the phylogenetic signal of parallel origins would be misleading, as the populations that are parapatric appear as each other's closest relatives. Conversely, when the *gsi* ratio is large and negative, we can be confident that the signal of a single origin represents the true species tree.

#### S4 Methods. *SLiM2* code for simulations

```
initialize() {
	if (exists("slimgui")) {
		defineConstant("seed", 1);
		defineConstant("mu", 1e-7);
		defineConstant("r", 1e-8);

		defineConstant("N", 100);
		defineConstant("t1", 10000);
		defineConstant("t2", 5000);
		defineConstant("mig", 0);
		defineConstant("tSec", 0.1);
		defineConstant("outPath", "~/workspace/PopGenSims/OriginScenarios");
	}

	setSeed(seed);
	initializeMutationRate(mu);
	initializeMutationType("m1", 0.5, "f", 0.0);
	initializeGenomicElementType("g1", m1, 1.0);
	initializeGenomicElement(g1, 0, 1e6 - 1);
	initializeRecombinationRate(r);
}

1 {

	// create ancestral population
	sim.addSubpop("p0", N);

	// schedule split and migration events based on parameter values
	t0 = 10*N;
	outGen = t0+t1+t2;

	sim.rescheduleScriptBlock(s1, start=t0+1, end=t0+1);
	sim.rescheduleScriptBlock(s2, start=t0+t1+1, end=t0+t1+1);
	sim.rescheduleScriptBlock(s4, start=outGen, end=outGen);

	if(mig == 0){
		sim.deregisterScriptBlock(s3);
	} else
	sim.rescheduleScriptBlock(s3, start=asInteger(t0+t1+1+round(t2*(1-tSec))), end=outGen);

}

s1 10 {

	// initial split of ecotypes here
	sim.addSubpopSplit("p10", N, p0);
	sim.addSubpopSplit("p100", N, p0);
	p0.setSubpopulationSize(10);
}

s2 20 {

	// output t1 vcf here
	outgroup = sample(p0.individuals, 1);
	ingroup = sapply(sim.subpopulations[sim.subpopulations != p0], "sample(applyValue.individuals, 30, replace =
F);");
	set = c(outgroup, ingroup);
	set.genomes.outputVCF(filePath = paste(c(outPath, "/", "t1.replicate-", seed, ".vcf"), sep=""), outputMultiallelics =
F);

	// subsequent split into populations here
	sim.addSubpopSplit("p11", N, p10);
	sim.addSubpopSplit("p101", N, p100);
}
```

```
s3 30 {

	// initialize migration here between parapatric divergent ecotypes
	p10.setMigrationRates(p100, mig);
	p100.setMigrationRates(p10, mig);
	p11.setMigrationRates(p101, mig);
	p101.setMigrationRates(p11, mig);
}

s4 40 late() {

	// output final vcf here
	outgroup = sample(p0.individuals, 1);
	ingroup = sapply(sim.subpopulations[sim.subpopulations != p0], "sample(applyValue.individuals, 30, replace =
F);");
	set = c(outgroup, ingroup);
	set.genomes.outputVCF(filePath = paste(c(outPath, "/", "t2.replicate-", seed, ".vcf"), sep=""), outputMultiallelics =
F);
}
```

#### S5 Methods. R code for the genealogical sorting index (*gsi*) calculations

```
# Code modified from: Ravinet, M. et al. The genomic landscape at a late stage of stickleback speciation: High genomic
divergence interspersed by small localized regions of introgression. PLoS Genetetocs 14, e1007358 (2018).

# ===== Load Dependencies =====
library(ape)
suppressMessages(library(vcfR))
library(geiger)
suppressMessages(library(adegenet))

# ===== Read Input Data =====
args <- commandArgs(trailingOnly = TRUE)
filename <- args[1]

#filename <- "~/Dropbox (OL)/OriginScenarios-results/N-1000.t1-10000.t2-10000.tSec-0.25.mig-1e-06/t2.replicate-1.vcf"
vcf <- read.vcfR(filename, verbose = FALSE)


# ===== Define gsi Function =====
gsi <- function(tr, grp){
  n <- length(grp) - 1
  # only consider internal nodes (tips get index 1:Ntip(tr))
  internal.nodes <- seq(Ntip(tr)+1, Ntip(tr) + Nnode(tr))
  # For each internal node, what are the descendant tips
  node.descendants <- lapply(internal.nodes, function(n) tips(tr, n))

  # ----- Jeff Groh 10 May 2019 -----
  # Previous code contained an error in the following lines:
    # Which nodes have descendants in the group being considered?
    # required <- sapply(descendants, function(x) any(grp %in% x) )
  # The problem with this is that it selects *all* nodes which contain *any*
  # members of the focal group. However, in the denominator for the gsi calculation,
  # we are only interested in summing the degrees of nodes which belong to the minimum
  # subtree that contains all members of the focal group. The code below fixes this by
  # selecting the correct set of nodes.

  # find root of mimimum subtree containing all members of focal group
  # how many tips of the focal group are descended from each node
  n.focal.members <- sapply(node.descendants, function(x){ length(which(grp %in% x)) })
  # how many total tips are descended from each node
  n.total.members <- sapply(node.descendants, function(x){ length(x) })
  # to be a root of the minimum subtree, a node must contain at least all members of the focal group
  candidate.subtree.roots <- which(n.focal.members >= length(grp))
  # Out of these, the node with the least number of total descendants will be the subtree root
  candidates.total.members <- n.total.members[candidate.subtree.roots]
  winner <- candidate.subtree.roots[which(candidates.total.members == min(candidates.total.members))]
  root.node <- internal.nodes[winner]
  # find all tips which descend from the min subtree root node
  subtree.tips <- tips(tr, root.node)
  # find all nodes whose descendents include any of those tips
  nodes.with.focal.descendants <- sapply(node.descendants, function(x)any(subtree.tips %in% x))
  # but with fewer descendants than that of the subtree root node
  node.depths <- node.depth(tr)[internal.nodes]
  root.depth <- node.depths[winner]
  # select required nodes for calculation
  required.nodes <- nodes.with.focal.descendants == TRUE & node.depths <= root.depth
  # ----- End Correction -----

  # How many connections to those nodes have? (tree is not necessarily
  # dichotomous)
  degree <- table(tr$edge)[ internal.nodes[required.nodes] ]
  #Ape takes one connection off the root node, so if d=2 then treat it as d=3
  # (no other nodes can have d=2)
  obs.gs <- n / (sum( degree - 2 ) + sum(degree==2))
  #minGS (basically same procedure but for  whole tree)
```

```
  degree.total <- table(tr$edge)[seq(Ntip(tr)+1, Ntip(tr) + Nnode(tr))]
  min.gs <- n / (sum( degree.total - 2 ) + sum(degree.total==2))
  gsi <- (obs.gs - min.gs) / (1 - min.gs)
  return(gsi)
}


# ===== Calculate GSI From Phylogenetic Tree =====
# Calculate gsi with respect to environment, that is, high gsi should reflect
# apparent monophyly of groups from the same location (multiple origins)
# rather than monophyly of true clades (single origin).
# Also calculate gsi for true clades so these can be compared.
# In vcf output from slim, individuals are organized sequentially as such:
# p0 (1 individual), p10, p11, p100, p101 (30 individuals each)
# where there is gene flow between p10 & p100 and also p11 & p101 (parapatric pairs).
# Create vectors of names of individuals that belong to these groups.
# This will be used as input for the gsi calculation.

all.inds <- colnames(vcf@gt)[-c(1:2)] # this vector starts with i1 (excluding outgroup)
loc1 <- all.inds[c(1:30,61:90)]
loc2 <- all.inds[c(31:60,91:120)]
clade1 <- all.inds[c(1:60)]
clade2 <- all.inds[c(61:120)]

# Calculate gsi for entire chromosome (1Mb)
gen <- as.matrix(vcfR2genlight(vcf))
tr <- root(nj(dist(gen)), outgroup = "i0", resolve.root = TRUE)

gsi.clade1 <- gsi(tr, clade1)
gsi.clade2 <- gsi(tr, clade2)
gsi.loc1 <- gsi(tr, loc1)
gsi.loc2 <- gsi(tr, loc2)


# ===== Output GSI Values =====
cat(paste(c(gsi.clade1, gsi.clade2, gsi.loc1, gsi.loc2), sep="\t"))
cat("\n")
```

#### Supplementary material References


Cummings MP, Neel MC, Shaw KL. 2008. A genealogical approach to quantifying lineage divergence. *Evolution* 62:2411–2422.

Haller BC, Messer PW. 2017. SLiM 2: Flexible, interactive forward genetic simulations. *Mol. Biol. Evol.* 34:230–240.

Paradis E, Claude J, Strimmer K. 2004. APE: Analyses of phylogenetics and evolution in R language. *Bioinformatics* 20:289–290.

Popescu AA, Huber KT, Paradis E. 2012. ape 3.0: New tools for distance-based phylogenetics and evolutionary analysis in R. *Bioinformatics* 28:1536–1537.
